# scMitoMut for calling mitochondrial lineage–related mutations in single cells

**DOI:** 10.1101/2024.08.03.606286

**Authors:** Wenjie Sun, Daphne van Ginneken, Leïla Perié

**Affiliations:** Institut Curie, Université PSL, Sorbonne Université, CNRS UMR168, Physique des Cellules et Cancer, 75005, Paris, France

**Keywords:** mitochondrial mutation, lineage-tracing, single-cell sequencing

## Abstract

Tracing cell lineages has become a valuable tool for studying biological processes. Among the available tools for human data, mitochondria DNA (mtDNA) has a high potential due to its ability to be used in conjunction with single-cell chromatin accessibility data, giving access to the cell phenotype. Nonetheless, the existing mutation calling tools are ill-equipped to deal with the polyploid nature of the mtDNA and lack a robust statistical framework. Here we introduce scMitoMut, an innovative R package that leverages statistical methodologies to accurately identify mitochondrial lineage related mutations at the single-cell level. scMitoMut assigns a mutation quality q-value based on beta-binomial distribution to each mutation at each locus within individual cells, ensuring higher sensitivity and precision of lineage related mutation calling in comparison to current methodologies. We tested scMitoMut using single-cell DNA sequencing, scATAC sequencing and 10× Genomics single cell multiome datasets. Using a single-cell DNA sequencing dataset from a mixed population of cell lines, scMitoMut demonstrated superior sensitivity in identifying small proportion of cancer cell lines compared to existing methods. In a human colorectal cancer scATAC dataset, scMitoMut identified more mutations than state-of-the-art methods. Applied to 10× Genomics multiome datasets, scMitoMut effectively measured the lineage distance in cells from blood or brain tissues. Thus, the scMitoMut is a free available (https://www.bioconductor.org/packages/devel/bioc/html/scMitoMut.html.), well-engineered toolkit for mtDNA mutation calling with high memory and CPU efficiency. Consequently, it will significantly advance the application of single-cell sequencing, facilitating the precise delineation of mitochondrial mutations for lineage tracing purposes in development, tumor and stem cell biology.

## Introduction

Cellular DNA barcode lineage tracing uses heritable DNA sequences as markers to track descendants from the same ancestral cells and has become a valuable tool for studying biological processes [1,2]. Cellular barcoding in humans uses retrospective methods based on genomic mutations. Using somatic genetic mutations as the markers with single-cell read-outs of genome sequencing has provided valuable insight into numerous biological fields, including development, aging and the onset of cancer [3–6]. However, this method is costly and fails to concurrently capture omics data related to cellular phenotypes, such as transcriptomics and epigenomics.

As an alternative, mutations in the mitochondrial DNA (mtDNA) have been used [7–9]. This has numerous advantages over somatic mutation–based cellular barcoding methods, including: the short size of mitochondrial DNA (mtDNA) (about 16,600 base pairs); the presence of multiple mtDNA copies (of 100s to 1000s in a blood cell [10]); and the much higher mutation rate of mtDNA (which is 100-to 1000-times that of nuclear genomic sequences) [11,12]. Further, as mtDNA lacks chromatin, it can be detected with a single-cell transposase-accessible chromatin sequencing (scATAC-seq) technology [8,9]. scATAC-seq can be used in conjunction with single-cell multiome sequencing to provides cellular phenotype information via transcriptomes or via chromatin accessibility data. However, the unique properties of mtDNA make it difficult to use methods designed for calling single-cell nuclear DNA mutations [13,14]. Additionally, searching for lineage-related mutations is not equivalent to searching for somatic mutations, as not all mtDNA mutations are lineage related [15]. Thus, better bioinformatics tools are necessary to analyse single-cell somatic mutations in mtDNA.

Current methods use the allele frequency of a single-nucleotide variant, and the number of cells containing the variant, to identify lineage informative mutations [16,17]. However, these tools then use a threshold based on the variant allele frequency (VAF) to call mutations in a single cell, yet VAF is affected by sequencing depths and noise [18], which could compromise sensitivity and specificity. To address this, statistical models such as beta-binomial distribution (BBD) have been proposed for scRNA-seq or bulk variant calling [19,20], but this has not yet been proposed for analysing mtDNA mutations.

In this work, we have developed a framework to call mtDNA mutations in single cells based on BBD. Testing this method using data with ground-truth and more complex data, we find that it has a better sensitivity as compared to existing methods. Notably, as this method gives a confidence score for each mutation in each cell, it provides an improved criteria for selecting lineage informative mutations for lineage analysis. We also have developed an R package (called scMitoMut, for single-cell mitochondrial mutation) that is available in Bioconductor, which uses H5F files to store process data (thereby reducing RAM usage) and written in C++ (reducing CPU consumption), which can be used on personal computers. The implementation of the BBD model and scMitoMut in R will serve as a valuable tool for enhancing mitochondrial mutation analyses and for using mitochondrial mutations in lineage tracing and mitochondrial genetic studies.

## Results

### Applying beta-binomial distribution to call lineage–informative mitochondrial mutations

We applied a hypothesis-testing framework to call mutations, which can be seen as a task to decide whether the observed read counts are generated by technical noise (e.g., PCR errors, sequencing errors, sample contamination or other unknown factors). After fitting a statistical distribution to the data, the framework compares the observed data to the fitted distribution, to evaluate the probability that the observed read counts are derived from noise. If this is unlikely, we reject the null hypothesis and accept the alternative hypothesis, thus calling a mutation. Using the probability value, we can quantify the fidelity of a cell containing a specific mutation by considering the magnitude of the noise **(Fig. 1b)**.

**Figure 1.**
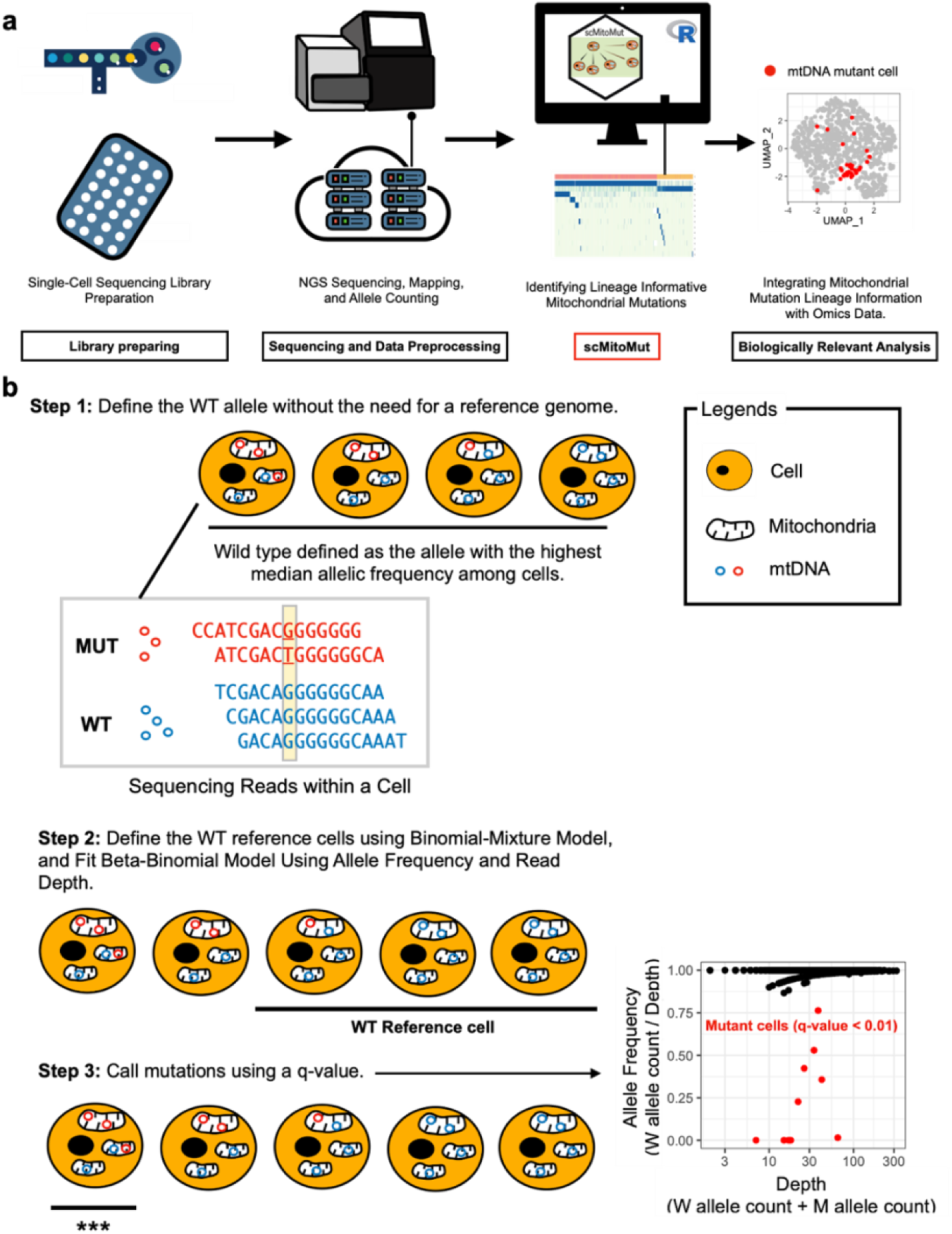
Overview of the scMitoMut package (a) After single-cell library preparation, sequencing data are pre-processed to generate the allele counting matrix generally using high performance computers or cloud computers. scMitoMut is then used to identify high-confidence single-cell mtDNA mutations using a personal computer. Finally, the mtDNA mutation results from scMitoMut are combined with cell phenotypes data for biological analysis. (b) Step 1 of scMitoMut consists of identifying the wild-type (WT) allele for each locus without the need of a reference genome. In brief, each cell contains multiple mitochondria, each with multiple mtDNA molecules (represented by circles). The read sequences obtained from the first cell, which has two alleles (in blue and red), are displayed at the bottom. Of all the cells, the blue allele has a higher median frequency (2/5, 2/4, 3/4, and 4/4 in the four example cells, respectively) than the red allele for the locus highlighted in yellow in the example sequences. This is defined as the WT allele. Step 2 consists of defining the WT reference cells using a binomial-mixture model classifier; the parameters of the beta-binomial model are then fitted to these cells for each locus using the allele frequency and sequencing depth reads count. Step 3 uses the beta-binomial model fitted in step 2 to call mutations for each locus in each cell. For a given allele frequency and sequencing depth reads count, the confidence score of the presence of a mutation (q-score) is calculated from the beta-binomial model, taking into account multiple comparisons using the false discovery rate (FDR). *** indicates cells with a q-value below a specified threshold are identified as mutant cells. The scatter plot on the right displays an example, in which allele frequency is plotted on the y-axis and depth on the x-axis. Each dot represents a cell, with q-values<0.01 in red, and other cells, in black.

As an input, the framework needs an allele frequency matrix per cell per locus. As a first step, we defined the wild-type (WT) allele. In contrast to previous methods that defined the reference genome base as the WT, we now defined the majority base in most cells as the WT, removing the effect of potential individual polymorphism compared to the reference genome. Once the WT allele identified, we defined the mutant reads as the total number of reads per locus minus the number of WT reads at the same locus for each cell. As in normal-depth mtDNA sequencing results, two mutations are rarely observed at the same site; this allowed us to reduce the statistical test needed for each cell, thereby increasing the statistical power with multi-test type I error correction **(Fig. 1b, Supplementary Fig. 1a,b**).

To select the reference WT cells for identifying mutations in subsequent steps, we classified cells into WT or mutant by fitting a binomial-mixture distribution to the mtDNA reads and computed the probability of specific cells belonging to the WT subset. Using this probability, we selected a cell subset assumed to have no mutations (by default, FDR > 0.05) **(Fig. 1b)**.

In the final step, we fitted the BBD to each locus using the WT cell subset **(Supplementary Fig. 1a)**. The fitted BBD model was then used as the null distribution to compute the probability of observing a specific WT allele, in a given cell and locus **(Supplementary Fig. 1b)**. To reduce the risk of calling a mutation due to multiple testing, the adjusted *p*-value, referred to as the mutation q-value, served as a quantitative measure of confidence in calling a mutation in a given cell and locus **(Fig. 1b, Supplementary Fig. 1b)**.

Overall, this statistical framework using BBD allows to call mtDNA mutations in single cells based on its q-value.

### Implementation of the scMitoMut package

We created an R package that we termed scMitoMut (freely available in Bioconcutor) that includes functions for data importing, mutation calling, data export and visualization **(Fig. 2a)**. In the analysis pipeline, the mapped BAM file is used to extract allele frequency using CellSNP-lite [21] or mtGATK [22]. Using the read count for the four bases (A, T, C and G) per cell per locus as the allele frequency, the allele frequency matrix is used by scMitoMut as an input for calling the lineage informative mutations per cell per locus. These lineage-informative mutations can then be jointly analysed with single-cell omics data **(Fig. 2a)**.

**Figure 2.**
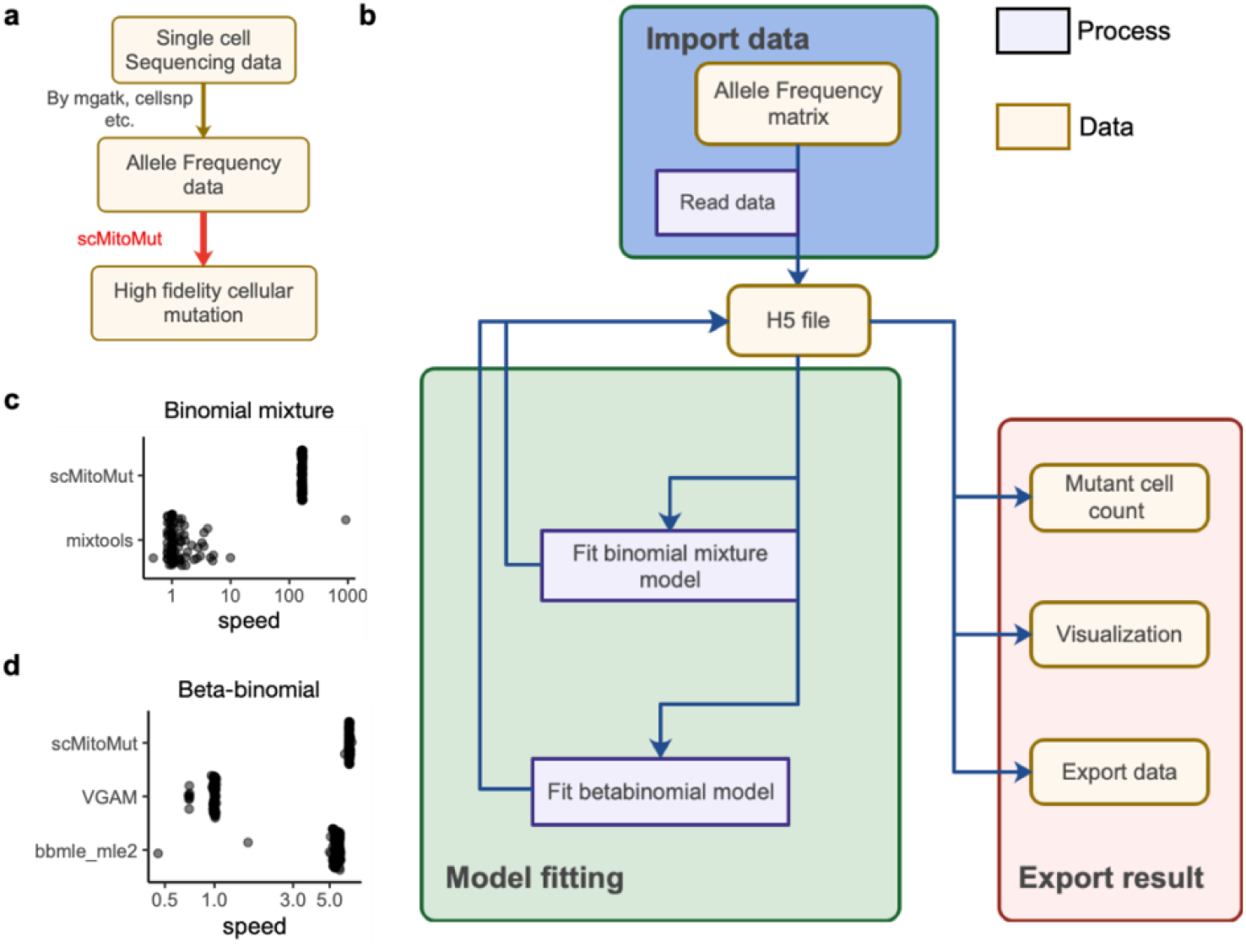
Tool chain and implementation of scMitoMut (a) Tool chain of using scMitoMut to call mtDNA mutations. The boxes represent the data and are linked by arrows representing the data operations. Tools used for the data operations are listed next to the arrows, with those conducted by scMitoMut highlighted in red. (b) The diagram illustrates the organization of the scMitoMut R package. The yellow box represents data connected by arrows symbolizing data operations. These operations describe the data operation processes (purple box). Rounded corner frames are used to emphasize groups of data and processes, showcasing the three main function in scMitoMut: data import (blue), model fitting (green), and result export (pink). All three functions interact with HDF5 files data for reading and writing raw and intermediate data stored on the hard disk. (c) Relative speed of each run divided by the median run time of mixtools for the R packages mixtools and scMitoMut. Each point is a run, with 100 runs tested per tool with the following parameters of the binomial mixture model: θ=0.5; π1 = 0.992; π2 = 1; n follows a log-normal distribution with log-mean = 2 and log-SD = 1 rounded up to the nearest non-negative integer; sample size is 5000. (d) Relative speed of each run divided by the median run time of VGAM for the R packages VGAM, bbmle, and scMitoMut. Each point is a run, with 100 runs tested per tool with 5000 observations and the following parameters of the beta-binomial model: mean probability parameter θ of 0.5 and a super-dispersion parameter ϕ of 20.

As a typical input matrix for scMitoMut included tens of thousands of cells, each of which has thousands of alleles and associated q-value, scMitoMut keeps all the raw input and intermediate output on a hard disk by using an HD5F database associated with an R object and functions for importing data, model fitting and exporting results. Thus, by reducing the RAM usage, scMitoMut can run on a personal computer and is able to analyse tens of thousands of cells without slowing down the system **(Fig. 2b)**.

We also boosted the mutation calling process by implementing expectation maximization and maximum likelihood estimation algorithms in C++ separately for fitting the BBD and binomial-mixture distributions. This significantly increased the speed of binomial-mixture fitting, achieving a maximum ∼170× improvement over the existing R function in the mixtools package [23] with the same fitting accuracy **(Fig. 2c, Supplementary Fig. 2a–c)**. Our BBD fitting achieved a fitting speed that was 6× faster than the VGAM package [24] with the same accuracy, and around 1.5× faster than the bbmle package with better accuracy **(Fig. 2d, Supplementary Fig. 3a–c)**. In scMitoMut, the fitting process can also run in parallel to leverage the multi-core capabilities of CPUs.

The scMitoMut is a well-engineered toolkit for mtDNA mutation calling with high memory and CPU efficiency that is freely accessible for researcher willing to study mtDNA mutation for lineage tracing.

### scMitoMut is more precise at identifying small clones from a scDNA-seq dataset of mixed cell lines

To compare the ability of scMitoMut to detect mutations as compared to other methods, we first tested the performance of scMitoMut using an in vitro experimental dataset with ground truth. We used a mix of BJ cells (human skin fibroblasts) and MKN45 cells (human gastric cancer cells) at a ratio of 99:1. MKN45 cells have copy number variation (CNV) but BJ cells do not, allowing us to distinguish the two types of cells. We identified 11 MKN45 cells and 1045 BJ cells, which served as ground truth for later benchmarking for mtDNA mutation calling **(Fig. 3a)**. The mtDNA sequencing depth distribution was relatively homogeneous over the loci, with a median distribution of around 10 **(Supplementary Fig. 4a)**. We then selected the cells with median distribution > 5, giving a final set of 962 cells (of which 11 MKN45 cells). This set was used to test the performance of scMitoMut with different parameters.

**Figure 3.**
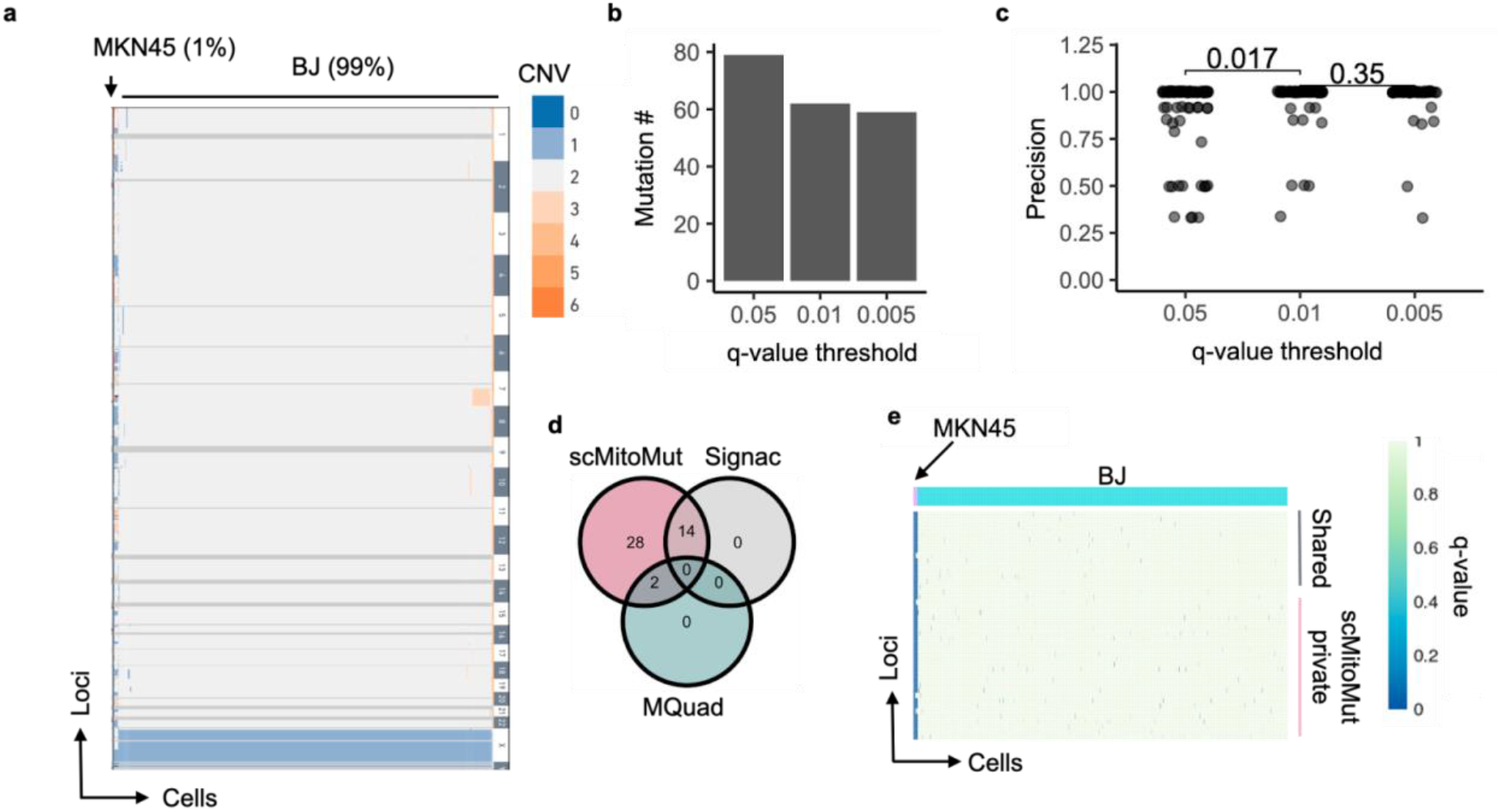
Benchmarking mtDNA mutation calling on a mixed–cell line dataset (a) Heatmap of the copy number variation (CNV) from single-cell genome sequencing data of a mixture of 11 MKN45 cells and 1045 BJ cells from a 10× Genomics demo dataset. Each row is a CNV segment, and each column is a cell. The color within the heatmap represents the CNV variation according to the color key on the right side of the heatmap, whereby the dark grey color indicates missing data, and light grey indicates diploidy. The cell types with and their percentages in the mixture are indicated at the top of the heatmap. (b) Number of mtDNA mutations present in MKN45 cells for various quality thresholds (q-values of 0.05, 0.01, and 0.005). The mutation was counted if its q-value was below the corresponding q-value in at least in one MKN45 cell. (c) Mutation calling precision for various q-value thresholds. The mutation precision was calculated as the ratio of mutant MKN45 to total mutant cells for each mutation. Each dot represents a mutation; p-values were calculated using the two-sided Wilcoxon test. The n is 79, 62 and 59 for q-value threshold of 0.05, 0.01 and 0.005 respectively. (d) Number of mtDNA mutations that were called in scMitoMut, Signac or MQuad analysis. Only mutations present in at least one MKN45 cell were counted. (e) Heatmap displaying the mutation quality q-values of scMitoMut. Each row represents a locus, and each column represents a cell. The darker the blue color, the lower the q-value, indicating higher confidence in the mutation, while the white indicates missing values. MKN45 and BJ cells were annotated in pink and blue above the heatmap. The mutations only detected by scMitoMut (28 total) are highlighted in pink, while shared mutations (16 total: 2 with MQuad, and 14 with Signac) are indicated in gray.

To determine how strictly we should call a mutation in a cell, we tested the mutation quality q-value (FDR adjusted *p*-value) thresholds of 0.001, 0.01 and 0.5. Increasing the q-value threshold led to a decrease in the number of observed mutations **(Fig. 3b)**. We calculated the precision for each mutation, defined as the percentage of MKN45 mutant cell numbers versus total mutant cell numbers. The majority of mutations exhibited a precision rate of 100%, with a statistically significant enhancement in precision observed when comparing q-value thresholds of 0.01 and 0.05 **(Fig. 3c)**. We saw no statistically significant improvement of precision when we used more stringent thresholds (from 0.01 to 0.005) **(Fig. 3c)**. When aggregating all mutations, the overall precision was 95.1% using a q-value threshold of 0.05, 98% with a q-value of 0.01, and 98.6% with a q-value of 0.005 **(Supplementary Fig. 4b)**. Notably, changing the q-value threshold did not statistically change the number of cells detected per mutation **(Supplementary Fig. 4c)**. Thus, scMitoMut precisely called mutations per cell, and a q-value of 0.01 can be used as default for calling a mutation in a cell.

To determine the minimum number of cells required to call a mutation, we tested the impact of different thresholds of cells per mutation on the precision and the numbers of lineage informative mutations. We observed no statistically significant differences in precision when increasing the minimum number of cells per mutation, despite an expected decrease in the total number of mutations called and an increase in clone size (number of cells per mutations) **(Supplementary Fig. 4d–g)**. Thus, the precision of the mutation calling results was controlled by the q-value alone. In the scMitoMut package, we therefore defined the default parameters as a minimum count of 5 mutant cells and a pre-determined q-value threshold of 0.01; note however that these values can be adjusted for different biological questions.

We next compared scMitoMut with the two state-of-the-art (SOTA) methods, Signac and MQuad [8,16,17]. Calling the mtDNA mutations in MKN45 cells using Signac, MQuad or scMitoMut (parameters are shown in **Supplementary Fig. 5a)**, we identified a total of 44 mtDNA mutations in MKN45 cells, of which scMitoMut detected all 44, Signac detected 14 and MQuad detected 2 **(Fig. 3d)**. The mutations identified by scMitoMut alone have the same qualities as the mutations also identified by Signac and MQuad. **(Fig. 3e, Supplementary Fig. 5b)**. Employing the previously defined precision (i.e., the proportion of MKN45 mutant cells to the total number of mutant cells), we found no statistically significant differences between mutations detected by scMitoMut and Signac **(Supplementary Fig. 5c)**. However, the median clone size of the Signac result is 10.5, which is statistically smaller than that of the scMitoMut result, of 11.0 **(Supplementary Fig. 6D)**. Thus, given that there are 11 MKN45 cells in this dataset, scMitoMut had better recall.

### scMitoMut package detects more mutations than other SOTA methods in scATAC-seq data from human colorectal cancer cells

We next tested the scMitoMut and other SOTA methods using a human colorectal cancer (CRC) scATAC-seq dataset with enriched mtDNA [22], which consists of epithelial-derived CRC cells and blood cells **(Supplementary Fig. 6a)**. We compared the lineage mutation identified by scMitoMut with the ones from MQuad and Signac. We used the default parameters for Signac and ScMitoMut (parameters are shown in **Supplementary Fig. 6b)** and the ones from the original publication for MQuad [17]. Of note, the original work used less stringent parameters for MQuad than the default settings or than those used in the two other methods [17]. Of the 14 mutations identified by scMitoMut, Signac and MQuad both identified nine mutations, whereby four mutations were identified only by scMitoMut and one only by MQuad (no specific mutations were identified by Signac) **(Fig. 4a, b)**. Thus, scMitoMut allowed us to identify more mutations, even as compared to MQuad, which used a less-stringent mutation filtering threshold. The MQuad-specific mutation was not identified by scMitoMut due to its clone size of four, which was below the threshold of 5 **(Supplementary Fig. 6b)**, but it would have been identified if the same threshold had been applied as in the original publication using MQuad [17]. Two of the scMitoMut-specific mutations, chrM.200 and chrM.310, were filtered by Signac by the VMR and strand concordance threshold, respectively **(Supplementary Fig. 6d,e,f)**.

**Figure 4.**
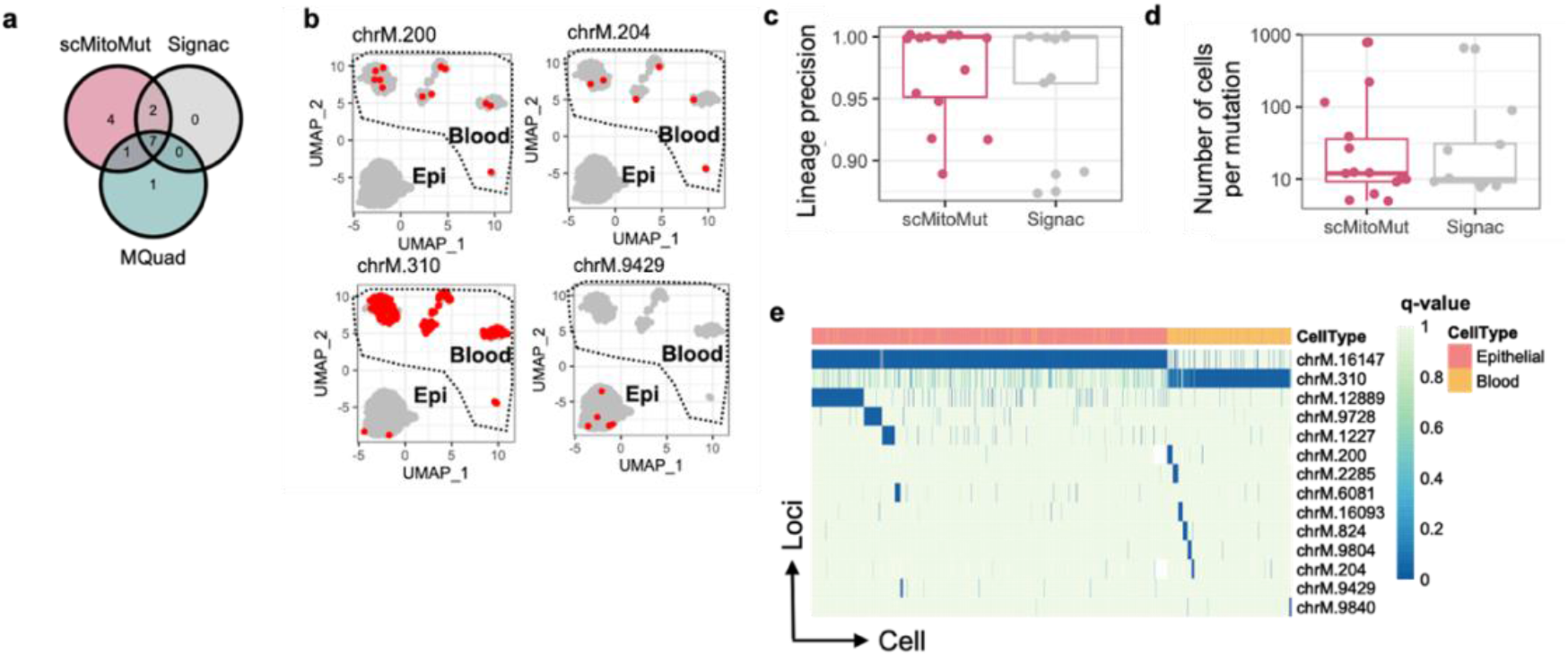
The beta-binomial model provides better lineage precision for distinguishing epithelial and blood lineages in the CRC dataset (a) Number of mtDNA mutations detected by scMitoMut, Signac and MQuad in the CRC dataset from Lareau et al. 2021. (b) The UMAP projections of the scATACSeq data with the cells with a mutation at one of the 4 loci exclusively detected by the scMitoMut. In each UMAP, the CRC and blood cell clusters are annotated. Each dot represents a cell, with red dots indicating high-confidence mutant cells called by scMitoMut with q-value < 0.01. (c) Mutation calling precision for each mutation in the scMitoMut or Signac results. The precision was calculated by dividing the count of cells in the predominant lineage (blood cells or cancer epithelial cells) by the total number of cells with a given mutation. n = 14 mutations in scMitoMut and n = 9 mutations in Signac. Center line, median; box limits, upper and lower quartiles; whiskers, 1.5× interquartile range; points, outliers are displayed. Two-sided Wilcoxon test was used to compare precision between the two methods (p = 0.89). (d) Number of mutant cells identified by scMitoMut or Signac. Each dot represents a mutation, n = 14 mutations in scMitoMut results and n = 9 mutations in Signac results. Center line, median; box limits, upper and lower quartiles; whiskers, 1.5× interquartile range; points, outliers are displayed. Two-sided Wilcoxon test to compare both methods (p = 0.92). (e) The mutation heatmap displays the scMitoMut mutation q-value per cell (columns) and per locus (rows). The deeper blue within the heatmap represents the smaller mutation q-value that correspond to higher confidence in the calling, while white indicates missing values. An annotation bar above the heatmap indicates the cell lineage is annotated above the heatmap, with cancer epithelial cells in red and blood cells in orange.

We then evaluated the lineage precision of mutations within a particular lineage of CRC and blood cells. Epithelial and blood cells diverge early in development, originating from the endoderm [25] and mesoderm [26], respectively, and should not share many mutations. Thus, defining lineage precision as the ratio of mutant cells in the dominant lineage of cancer epithelial or blood cells as compared to all mutant cells, we expect a 100% lineage precision if the mutations were exclusively present in one lineage. The lineage precision was similar between scMitoMut and Signac **(Fig. 4c)**, indicating that the detected mutations by scMitoMut were informative for lineage identification. While scMitoMut found more mutations, the number of cells detected per mutations was similar between scMitoMut and Signac **(Fig. 4d)**, suggesting that scMitoMut detected mutations with a similar cell abundance as previous methods. All four of the scMitoMut-specific mutations are lineage informative mutations that are restricted to one lineage of cancer epithelial or blood cells **(Fig. 4b)**.

In the mutation q-value heatmap generated from scMitoMut results, the mutations chrM.16147 and chrM.310 are predominantly confidently detected (with small q-value) in CRC and blood cells, respectively **(Fig. 4e)**. Other mutations appeared to be clustered in regions on either chrM.16147 or chrM.310, indicating that the chrM.16147 and chrM.310 mutations appeared earlier **(Fig. 4e)**. The allele frequency heatmap is consistent with the q-value heatmap, but with a lower signal-to-noise ratio between mutant cells and WT cells; this is especially seen for the later mutations clustered on chrM.16147 and chrM.310 **(Supplementary Fig. 6b)**, demonstrating the added value of using the q-value to call and visualize lineage informative mutations.

### mtDNA mutations identify lineage branching in two 10× multiome human datasets

After determining that scMitoMut exhibited higher sensitivity and specificity as compared to the SOTA methods Signac and MQuad in single-cell, whole-genome sequencing and mtDNA-enriched scATACSeq datasets, we next tested scMitoMut on a dataset for which both the chromatin accessibility and the transcriptome were recovered from the same cells using the 10× Genomics multiome kit. We first analysed the 10× Genomics Inc single-nuclear multiome dataset from human peripheral blood mononuclear cells (PBMC). After analysing the transcriptomes of the cells, we divided cells into three categories: monocytes, T cells and B cells **(Fig. 5a)**. Even though single-nucleus sequencing was used, mtDNA reads can still be found in the scATAC-seq results with median depth of 3, which indicates that mtDNA could be detected (**Supplementary Fig. 7a)**. Thus, we included the cells with mean mtDNA depth of > 5, resulting in median mtDNA sequencing depth of 10 **(Supplementary Fig. 7b)**; this is comparable to the mtDNA profile of single-cell genome sequencing of a mixture of cells that we analysed before (see **Supplementary Fig. 4a)**. Finally, there were no noticeable differences in mtDNA sequencing depth for monocytes, T cells or B cells **(Supplementary Fig. 7c–e)**.

**Figure 5.**
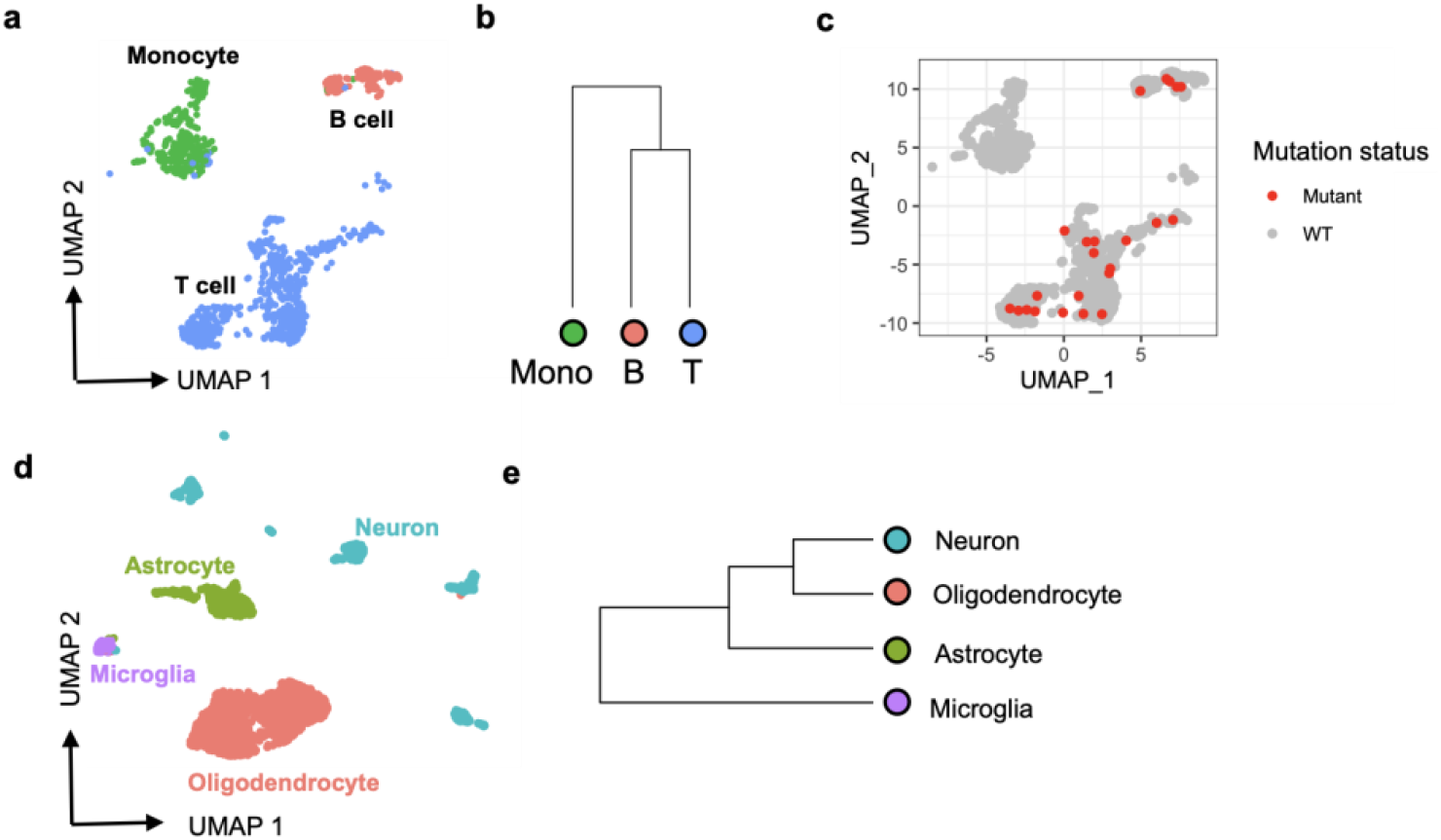
scMitoMut identifies lineage-informative mitochondrial mutations in the 10X multiome dataset. (a) UMAP visualization of the PBMC 10X multiome dataset with default parameters using Seurat. Each dot represents a cell, color-coded by cell type: red for B cells, blue for T cells, and green for monocytes. (b) Hierarchical clustering of monocytes (shown in green), B cells (in red), and T cells (in blue), based on mutant cell counts normalized by total mutant count per cell type cluster, using Euclidean distance and complete linkage clustering. (c) UMAP visualization of locus chrM.16499. Each dot represents a cell, and red dots indicate mutant cells (q-value < 0.01) identified using scMitoMut. (d) UMAP vizualization of brain data from 10× multiome dataset. Each dot represents a cell; oligodendrocytes are shown in red, microglia in purple), neurons in green and astrocytes in blue. (e) Hierarchical clustering analysis of four brain cell types conducted by the method described in (b)

We then analysed mtDNA mutations in single-cell ATAC-seq data using scMitoMut; this identified a total of 17 mutations based on a minimum mutant cell number of > 10 and a mutation quality value q-value threshold of 0.01 **(Supplementary Fig. 8a,b)**. In comparison, MQuad and Signac detected fewer mutations using all their default parameters except the minimum mutant cell number requirement of > 10 **(Supplementary Fig. 8a,b)**. This demonstrated that scMitoMut had a higher sensitivity also for multiome data.

Taking advantage of this higher sensitivity, we conducted hierarchical clustering on the three cell clusters based on the normalized mutant cell count per cell type per mutation. Of note, Signac lacked sufficient mutations for tree reconstruction, and MQuad does not incorporate tree reconstruction. The results indicated that the two lymphoid cell types (T and B cells) clustered together and were separated from the monocytes **(Fig. 5b)**. This aligns with our current understanding of blood cell development, as T and B cells originate from the lymphoid progenitor, while monocytes derive from the myeloid progenitor. It also demonstrated that the increase sensitivity of scMitoMut allows cells to be correctly assigned to their lineage of origin.

Studying the central nervous system development in mice has revealed that microglia are derived from yolk sac cells in embryo, while neurons, astrocytes and oligodendrocytes are derived from embryonic neural tube in embryos [21,27]. To explore the lineage distance between cells in the human central nervous system, we analysed brain cell 10× multiomics data, which included 3,000 brain cells, with scATAC-seq and scRNA-seq. We annotated neurons, astrocytes, oligodendrocytes and microglia using known cell type markers **(Fig. 5e, Supplementary Fig. 9c)**. Using the same parameters as applied in the above 10× Genomics multiome PBMC dataset, scMitoMut identified 14 mutations, and MQuad identified four mutations. However, none of these mutations passed the threshold set by Signac **(Supplementary Fig. 9d)**. Using the 14 mutations identified by scMitoMut, we calculated the mutation frequency for each cell type and conducted hierarchical clustering based on these frequencies. This analysis revealed that human microglia are less closely related to neurons and oligodendrocytes, similar to the known patterns of cell development in the central nervous system of mice **(Fig. 5f)**.

Thus, by applying scMitoMut to two human datasets, we demonstrated that it can be used to assess lineage relationship between cells, which is of a particular importance for understanding human lineage, as most of it is unknown.

## Discussion

Here we present scMitoMut, an R package that we developed to facilitate the identification and analysis of mitochondrial lineage informative DNA mutations at a single-cell resolution. Within the scMitoMut package, we implemented a BBD-based statistical framework to call the mtDNA lineage informative mutation. Using this statistical framework, scMitoMut provides a q-value for the confidence of the presence of the mutation at each locus that is used to call if the mutation is present in a single cell. This approach allows the user to select lineage-informative mutations based on the tools from statistical testing (in contrast to thresholds based on reads, as in other SOTA methods); likewise, it demonstrated heightened sensitivity in mutation detection as compared to the other SOTA methods.

In practice, scMitoMut includes an efficient implementation using an HDF5 file database for high-speed data access and reduced runtime memory usage. To enhance running speed, we used Rcpp for distribution fitting and parallelised tasks with CPU multi-threading. This package will thus serve as an invaluable tool for enhancing mitochondrial mutation analysis as well as for promoting the use of mitochondrial mutation in lineage tracing and mitochondrial genetic studies in large dataset, such as the atlas dataset [28]. Compared to other SOTA methods [8,16,17], scMitoMut uses fewer parameters, of only the quality q-value and clone size parameters, thus providing an easier-to-use tool.

The flexibility of the beta-binomial model makes it more robust at learning the noise distribution if there is still be a small proportion of mutant cells in the reference cells used for fitting. However, the presence of a high proportion of mutant cells can cause overfitting of the beta-binomial and thus weaken the ability of the beta-binomial model to detect mutant cells. To overcome this issue, we used a binomial-mixture classifier for unsupervised classification.

By analysing single-cell mitochondrial mutations of known lineages, we found that the precision of mutations is related to the q-value threshold rather than to the threshold of the number of cells with a given mutation. However, in practice, the total number of false-positive mutations is a product of both the mutation precision and the total number of detected mutations. Lowering the threshold of the mutant cell number leads to a higher detection of mutant cells, which increases the total number of false-positive mutant cells. We recommend adjusting the threshold of the mutant cell number based on usage purposes.

Mitochondrial somatic mutations have recently emerged as an optimizing lineage genetic marker, which hold great potential in human lineage tracing [8,29]. However, some issues are still seen as hindrances, including how the high mutation occurrence and the fast turnover rate of mutations in mtDNA affect the lineage information [15], and how the RNA editing issues affect calling lineage informative mutation [8]. While more work is required to investigate the impact of these issues on the use of scMitoMut, we currently recommended using DNA sequencing (e.g., scATAC-seq) as the input value for scMitoMut to avoid RNA editing. In this study, we focused on identifying mutations with lineage information; how to best use these mutations for lineage reconstruction should now be investigated.

In summary, in the newly-developed scMitoMut package, we have established a robust statistical framework for identifying lineage-informative mtDNA mutations, with an easy-to-use interface in R. This tool enhances the utilization of mtDNA somatic mutations to trace lineages.

## Methods

### Applying beta-binomial distribution to call lineage informative mitochondrial mutations

#### Defining the wild-type allele frequency

Single cell sequencing data provided allele counts per locus *j* = 1,2, …, *K* per cell *i* = 1,2, …, *S*. In the single nuclear variation case, for a specific locus and cell, the allele count *n* for four nucleotides are: *n*_*A,i,j*_, *n*_*T,i,j*_, *n*_*C,i,j*_, and *n*_*G,i,j*_. The sequencing depth *N*_*i,j*_ was defined as:

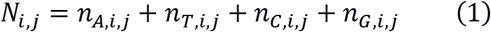

To define the wild-type allele, we used the dominant allele *W*_*j*_ for locus *j* over all cells:

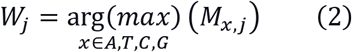

ere the *M*_*x,j*_ was the median count of allele *x* for locus *j* across all cells.

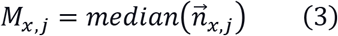

In the equation (3) the 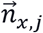 was the count of allele *x* at locus *j* across all cells. Specially:

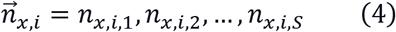

For a cell *i* and locus *j*, let *m*_*i,j*_ represent the allele count for the wild type allele *W*_*j*_, which was used to calculate wild type allele frequency *wAF*_*i,j*_:

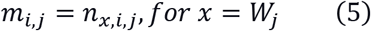

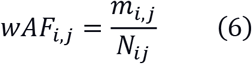

We used *wAF*_*i,j*_ as a proxy of the heteroplasmy level of the wild type allele at locus *j* of cell *i*.

Based on previous publications [9,17,22,22], we assumed that each locus was unlikely to have more than one mutation. Therefore, each locus could either be wild type or bear a single mutation, representing two possible states. These states were modeled using binomial related models. A mutation is identified for locus *j* in cell *i* when the allele frequency *wAF*_*i,j*_ is significantly lower.

### Fitting mitochondrial heteroplasmy sequencing results with beta-binomial distribution

Since mutation calling for each locus was independent of each other in our strategy, we explained the model fitting process for a single locus, omitting the subscript *j* for conciseness.

Single-cell sequencing can be modelled as a random sampling process of mtDNA (mitochondrial DNA) from a cell. Assuming the sampled mtDNA number is small compared to the total mtDNA copy number in a cell, the sampling process of sequencing mtDNA then corresponds to a series of Bernoulli trials. In this Bernoulli framework, sequencing the wild type allele is defined as success with the probability *π*, which correspond to the mtDNA heteroplasmy level of the cell. *π* is inferred from AF using the resulting successful trial number *m*_*i*_ (wild type allele count of cell *i*) and the total trial number *N*_*i*_ (sequencing depth of cell *i*). Let random variable *X* represents the wild type allele count, it then follows a binomial distribution when sequencing a cell with heteroplasmy level *π* and total allele count *N*_*i*_.

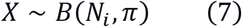

Mitochondrial turnover can create inter-cell heterogeneity, which create heterogenous mtDNA heteroplasmy *π* between cells. To account for these factors, we introduce the Beta-binomial model which allows *π* to be a random variable. In the Beta-binomial model, we consider the mtDNA heteroplasmy level as a continuous random variable *Π* ∈ [0,1], and is described by the Beta-distribution:

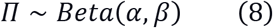

Let *X* be the wild type allele count and a random variable which follows a beta-binomial distribution (BBD):

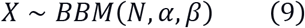

The BBD also can be described in this format:

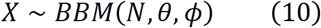

Where,

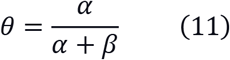

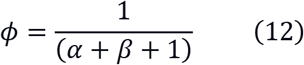

The parameter *θ* describes mean of *Π*, while the *ϕ* is the dispersion parameter for describing the variance. If the *ϕ* = 0, the BBD degrades into binomial distribution, with a probability parameter of *θ*. To avoid fitting the BBD on mutant cells (which will cause over-fitting), cells were clustered using binomial-mixture distribution to select the likely WT cells for model fitting. After fitting the beta-binomial model, the hypothesis test was used to get probability of a mutation using Beta-binomial PMF function:

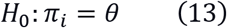

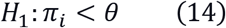

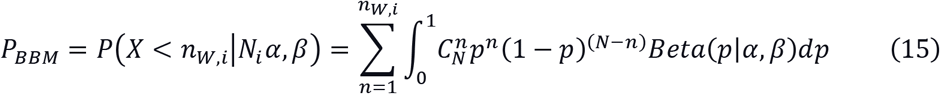

We got *P*_*BBM*_ per locus per cell. For each locus, we performed the false discovery rate (FDR) correction for multiple tests across multiple cells, and nominated the FDR value as q-value for evaluating the quality of lineage informative mutation.

### Clustering cells with binomial mixture model

We defined the WT allele heteroplasmy level in WT cell *π*_*W*_ and in mutant cell *π*_*M*_; and the percentage of WT cells *θ*_*W*_ and mutant cells *θ*_*M*_ in the population. Let the random variable *X* of the WT allele count follows a binomial mixture model:

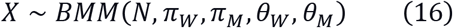

After fitting the model to the data, we calculate the probability of cell under investigation having a WT allele is calculated; this is denoted as *P*_*BMM*_.

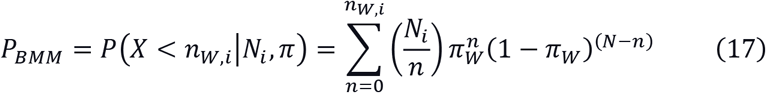

The value of the probability *P*_*BMM*_ is used to classify the cells to get potential WT cells for fitting the beta-binomial model. The likely WT cells are selected by binomial-mixture model with an FDR (false discovery rate) adjusted *P*_*BMM*_ value larger than 0.05.

### Rcpp-based distribution fitting algorithm reimplementation and benchmarking

We reimplemented the model fitting algorithm of beta-binomial mixture and binomial-mixture distribution using Rcpp in R [30]. This reimplementation was intended to improve CPU efficiency. The beta-binomial model was fitted using maximum likelihood estimation (MLE), which finds the parameters of the BBD that maximize the likelihood function that represents the probability of the observed data given the model parameters. The binomial-mixture distribution was fitted with expectation-maximization (EM) fitting, which is an iterative method to find MLE of parameters in models with latent variables with iterations of expectation (E) step and maximization (M) step. Detailed mathematical formulations, derivations, and additional implementation details can be found in the supplementary methods section.

We compared the speed of fitting binomial-mixture distribution between scMitoMut and the multmixEM function in the mixtools R package [23], by fitting simulated binomial mixture distribution with: two binomial distributions with *θ* = 0.5, *π*1 = 0.992 and *π*2 = 1; the *n* follows a log-normal distribution (log-mean = 2 and log-SD = 1) rounded up to the nearest non-negative integer. Each simulation includes 1000 observations. The simulation was done in R with a custom function using rlnorm and rbinom R random number generator. We benchmarked the running speed with the microbenchmark function from the microbenchmark R package with default parameters [31]. The accuracy of model fitting was tested using the simulations follow the parameter: *π*_1_ = 0.5, *π*_2_ = 1, *θ* = 0.01, 0.1, 0.2 *or* 0.4 and *n* following a log-normal distribution (log-mean=2, log-SD=1) rounded to the nearest non-negative integer; each simulation contains 5000 observations.

We benchmarked the model fitting speed of beta-binomial model between scMitoMut, vglm function from VGAM R package [24] and mle2 function from bbmle R package [32], by fitting the simulated data with: mean probability parameter *θ* of 0.5; super-dispersion parameter *ϕ* of 20; each simulation includes 5000 observations. The simulation was done by the “rbetabinom.ab” function from VGAM R package [24]. The mle2 function is a generated maximum likelihood fitter, we need to provide the target function which was the dbetabinom function from the emdbook R package [33]. The accuracy of the fitting was tested using a simulated beta-binomial distribution with parameters specified in rbetabinom.ab: mean parameter *θ* of 0.1, 0.5, 0.9, or 0.99; dispersion parameter *ϕ* of 20, 40, 80, 160; *n* adheres to a log-normal distribution with a log-mean of 2 and log-SD of 1, subsequently rounded up to the nearest non-negative integer; each simulation encompasses 5000 observations.

### Mixed–cell line single-cell genome sequencing

#### Identifying the MKN45 and BJ cells in the mixed cell line dataset using CNV

The dataset consists of single-cell genome sequencing data for a cell line–mixture containing 1% MKN45 cells and 99% BJ cells from the 10× Genomics demo. In the dataset, the MKN45 cells, a cancer cell line with chromosome instability, were distinguished from the normal BJ cell line using CNV analysis. A Loupe scDNA Browser 1.1.0 was used to analyse the “bj_mkn45_1pct_dloupe.dloupe” output from CellRanger DNA version 1.1.0, which was downloaded from 10× Genomics Demo dataset website. Using Loupe scDNA Browser, we generated CNV heatmap, and extracted cell clusters based on CNV. Based on CNV, we identified a MKN45 cell cluster with 11 cells (1%), which has high CNV variations, from 1045 cells. The cellular barcodes of MNK45 cells were exported from Loupe scDNA Browser and used in the future as ground truth of the MKN45 cell identity for benchmarking the mtDNA lineage informative mutation calling.

#### Calling mtDNA mutation using scMitoMut

The mgatk version 0.7.0 was used to analyse the Bam file to count mtDNA allele frequency. The mgatk ran in “tenx” mode options “--keep-duplicate” and no alignment-quality constraint “ -- alignment-quality -1”. The Bam file “bj_mkn45_1pct_possorted_bam.bam” was downloaded from 10X Genomics website together with the good quality cell list “bj_mkn45_1pct_per_cell_barcode.tsv”. The “coverage.txt.gz” file within the mgatk output was used to assess coverage distribution per locus for all cells. After selecting cells with an average mtDNA coverage greater than 5, scMitoMut calculated the q-value for mutation quality per cell per locus using the allele count matrix from mgatk.

We identified mutations with a q-value below a threshold per cell per locus, and required the number of mutant cells to meet a threshold. We assessed the specificity of mtDNA lineage-informative mutations using various q-value thresholds (0.05, 0.01 and 0.005) and mutant cell number thresholds (1, 5, and 9). The identified mutation number and mutation precision were evaluated using the mutations with at least one mutant MKN45 cell.

For each mutation, we defined the precision of mutation calling as the percentage of MKN45 cells. For a specific mutation *i*, the mutant cells number in MKN45 cells is *n*_*i*_ (*n*_*i*_ > 0) and *m*_*i*_ for total mutant cells. We defined the precision *y*_*i,pre*_ by dividing mutant MKN45 cell to the total mutant number:

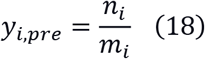

We calculated the overall precision *y*_*pre*_ by dividing the total number of mutant MKN45 cells by the total number of mutant cells after merging all identified mutations.

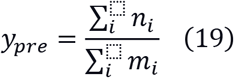

We used a q-value of 0.01 and minimum mutant cell number of 5, when comparing the mutation calling results with other SOTA methods.

#### Calling mtDNA mutation using Signac

The Signac R Package is designed for scATACSeq analysis, which also equipped mtDNA mutation calling function by providing the ReadMGATK, IdentifyVariants functions [16]. We used the ReadMGATK function to read in the mgatk output which was also used in scMitoMut mutation calling process.

For the Signac, we followed the vignette and default parameters: VMR (variant mean ratio) larger than 10^-2; strand accordance at least 0.65; mutant cells at least 5, where the mutant cells were defined by at least 2 mutant reads in both fwd and rev strand.

#### Calling mtDNA mutation using MQuad

We utilized CellSnip-lite Version 1.2.3 to process the bam file and generate the allele frequency matrix using the --countORPHAN option. The resulting files “tag.AD.mtx” and “tag.DP.mtx” were then inputted into MQuad Version 0.1.8 with a minimum DP of 5 specified (--minDP 5). Mutations were counted based on whether their DeltaBIC value exceeded the Kneel point, as indicated by the “PASS_KP” flag in the “BIC_params.csv” file. No minimum mutant cell number threshold was applied, making the filtering less stringent than the Signac or scMitoMut pipeline.

### Colorectal cancer scATACSeq mitochondrial mutation analysis

#### Cell type annotation

We used the colorectal cancer (CRC) scATAC-seq with mtDNA kept (mtscATAC-Seq) from Lareau et al. [22]; the scATAC-seq file can be download following the links in the Signac package’s (v1.9.0) vignettes (https://github.com/stuart-lab/signac/blob/1.9.0/vignettes/mito.Rmd).

We followed the instructions provided in the vignette to preprocess the scATACSeq data. We annotated open chromatin peaks with the EnsDb.Hsapiens.v75 gene. The high-quality cells fulfilling all the following conditions were retained: nCount_peaks > 1000, nCount_peaks < 50000, pct_reads_in_DNase > 40, blacklist_ratio < 0.05, TSS.enrichment > 3, nucleosome_signal < 4, and an average mitochondrial sequencing depth >= 10. We normalized peaks with the term frequency inverse document frequency (TFIDF) method. Peaks with a minimum of 10 counts are selected and subjected to singular value decomposition (SVD) for dimension reduction. Dimensions 2 to 50 were utilized to generate the UMAP and perform nearest neighbor-based clustering with LSI reduction, using a resolution of 0.5 and algorithm 3. Gene accessibility of TREM1, EPCAM, PTPRC, IL1RL1, GATA3, and KIT were employed to annotate cells into epithelial, basophil, myeloid, and T cell categories. These cell clusters were subsequently binarized into epithelial or blood cell categories.

#### Comparing the mutation calling between scMitoMut, and MQuad Signac

scMitoMut used the allele count matrix output from mgatk, which was downloaded using the link in the Signac vignette (https://github.com/stuart-lab/signac/blob/1.9.0/vignettes/mito.Rmd). The mutant cell was called by q-value < 0.01, then the mutations were filtered with a mutant cell number threshold of 5.

For Signac, we adhered to the guidelines provided in the Signac package (v1.9.0) vignette (https://github.com/stuart-lab/signac/blob/1.9.0/vignettes/mito.Rmd). Mutations are called by imposing the following thresholds: VMR (variant mean ratio) larger than 10^-2, strand accordance > 0.65, and mutant cells ≥5, where the mutant cells were defined by at least 2 mutant reads in both fwd and rev strand (cells_conf_detected >=5, strand_correlation >= 0.65, and vmr > 0.01).

The MQuad result was retrieved from Kwok et al. [17], https://github.com/aaronkwc/MQuad_paper_reproduced_results/blob/main/CRC_mtscATAC/data/CRC_mgatk.csv. They called all the mutations with deltaBIC < 0, which means an informative gain for binomial-mixture versus binomial model, without mutant cell number threshold.

For each mutation, we defined the precision of mutation calling as the consistency with cell types. For a specific mutation *i*, the mutant cells number in each lineage is *n*_*i*_ and *m*_*i*_ for epithelial and blood lineage respectively. We defined the precision *y*_*i*_ by dividing the dominant lineage to the total mutant number:

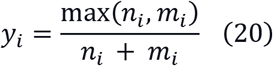

The precision *y*_*i*_ is a rational number in the range of [0.5, 1].

### PBMC 10× Genomics multiome data

#### Cell type annotation

The 10× Genomics multiome dataset pbmc_granulocyte_sorted_10k was downloaded from 10× Genomics website. We preprocessed the data according to the methods described in multiome data analysis vignette of Signac (https://github.com/stuart-lab/signac/blob/1.9.0/vignettes/pbmc_multiomic.Rmd). Briefly, we selected the high-quality cells fulfilling all of the following condition: nCount_ATAC < 100000, nCount_RNA < 25000, nCount_ATAC > 1000, nCount_RNA > 1000, nucleosome_signal < 2, TSS.enrichment > 1. With Seurat 4.4.0, the transcriptome data was normalized using SCT transform, followed by PCA and UMAP embedding. The reference annotation dataset used was from Hao et al. 2021 [34]. We annotated cells as CD14 Mono, CD16 Mono, CD4 Naive, CD4 TCM, CD4 TEM, CD8 Naive, CD8 TCM, CD8 TEM, Treg, CD4 CTL, gdT, MAIT, B memory, B intermediate, B naive. We then classified these cell types into three categories: mono, T or B.

#### Calling mtDNA mutation using scMitoMut, Signac and MQuad

The mgatk version 0.7.0 ran in “tenx” mode options “--keep-duplicate” and no alignment-quality constrain “ --alignment-quality -1”, with the Bam file “pbmc_granulocyte_sorted_10k_atac_possorted_bam.bam”, and QC-passed cell list from “filtered_feature_bc_matrix.tar.gz” file downloaded from 10× Genomics website. The “coverage.txt.gz” of mgatk output was used to evaluate the coverage distribution per locus for all cells.

For scMitoMut mutation calling, we utilized cells with an average mtDNA depth greater than 5. The mutation calling threshold was set at a q-value of less than 0.01 and required a minimum of 10 mutant cells.

For Signac, we applied the parameters: VMR (variant mean ratio) larger than 10^-2, strand accordance > 0.65, and mutant cells >= 10, where the mutant cells were defined by at least 2 mutant reads in both fwd and rev strand (cells_conf_detected >=5, strand_correlation >= 0.65, and vmr > 0.01).

For MQuad, the allele frequency matrix was prepared by CellSnp-lite Version 1.2.3 with options “--UMItag None --countORPHAN --exclFLAG UNMAP --chrom chrM”. Then, MQuad Version 0.1.8 was used with minimum DP of 5 specified (--minDP 5). Mutations were called based on whether their DeltaBIC value exceeded the Kneel point, as indicated by the “PASS_KP” flag in the “BIC_params.csv” file, and more than 10 mutant cells per mutation.

### Brain 10× Genomics Multiome data

#### Cell type annotation

The 10× Genomics multiome dataset human_brain_3k was downloaded from 10× Genomics website. We selected the high-quality cells fulfilling all of the following conditions were retained: nCount_ATAC < 100000, nCount_RNA < 25000, nCount_ATAC > 1000, nCount_RNA > 1000, nucleosome_signal < 2, TSS.enrichment > 1. The transcriptome data was normalized using SCT transform, followed by PCA and UMAP embedding (with 30 PCA dimensions). The cell clusters were identified using FindNeighbours and FindClusters function with default parameters. Then we used the gene expression markers to annotated the cell clustering with MAP2 and SYN1 for neuron [35]; GFAP and ALDH1L1 for astrocyte [36]; P2RY12 and CX3CR1 for microglia [37]; MBP for oligodendrocyte [38].

#### Calling mtDNA mutation using scMitoMut, Signac and MQuad

The mgatk version 0.7.0 ran in “tenx” mode options “--keep-duplicate” and no alignment-quality constrain “--alignment-quality -1” with the Bam file “human_brain_3k_atac_possorted_bam.bam” and QC-passed cell list in “filtered_feature_bc_matrix.tar.gz” downloaded from 10X Genomics website. Within the mgatk output, the coverage per locus per cell file “coverage.txt.gz” was used to evaluate the coverage distribution per locus for all cells.

We called the mtDNA mutation using scMitoMut with the cells with average mtDNA depth larger than 5. The thresholds were set to: q-value < 0.01 and mutant cell number at least of 10.

For Signac, we applied the parameters: VMR (variant mean ratio) larger than 10^−2^, strand accordance > 0.65, and mutant cells >= 10, where the mutant cells were defined by at least 2 mutant reads in both fwd and rev strand (cells_conf_detected >=5, strand_correlation >= 0.65, and vmr > 0.01).

For MQuad, the allele frequency matrix was prepared by CellSnp-lite Version 1.2.3 using the bam file with options “--UMItag None --countORPHAN --exclFLAG UNMAP --chrom chrM”. Then, MQuad Version 0.1.8 was used wih minimum DP of 5 specified (--minDP 5). Mutations were counted based on whether their DeltaBIC value exceeded the Kneel point, as indicated by the “PASS_KP” flag in the “BIC_params.csv” file, and more than 10 mutant cells per mutation.

## Statistics

### Statistical test

The two-sided Wilcoxon rank-sum test was used to compare continuous unpaired variables, while the two-sided Fisher exact test was applied to categorical data. The false discovery rate (FDR) was applied for multiple tests correction. The heatmap is sorted using the “memo sort” method adapted from https://gist.github.com/armish/564a65ab874a770e2c26.

### Clustering cell types using mtDNA mutation

Let *C* be the set of cell types, and *M* be the set of mutations. We defined *N*_*ij*_ as the number of cells of type *i ∈ C* that have mutation *j ∈ M*. For each mutation *j ∈ M*, we normalized the number of mutant cells per cell type. Define *T*_*j*_ as the total number of cells with mutation *j*:

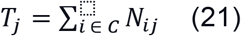

The normalized mutation frequency *F*_*ij*_ for cell type *i* and mutation *j* is given by:

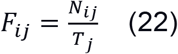

Using the normalized mutation frequency matrix *F*, we calculate the Euclidean distance between cell types based on their mutation frequencies followed by the hierarchical clustering using complete methods in R hclust function [39].

## Code availability

The code for all analyses in this study is available on GitHub (https://github.com/TeamPerie/scMitoMut_paper_SUN_et_al). The scMitoMut package is available on Bioconductor (https://bioconductor.org/packages/release/bioc/html/scMitoMut.html or https://doi.org/10.18129/B9.bioc.scMitoMut).

## Data availability

The 1k BJ dataset with 1% MKN45 spike-in for single-cell DNA sequencing was obtained from https://www.10xgenomics.com/cn/datasets/1-k-bj-with-1-percent-mkn-45-spike-in-1-standard-1-1-0. The human PBMC multiome sequencing data was downloaded from https://www.10xgenomics.com/cn/datasets/pbmc-from-a-healthy-donor-granulocytes-removed-through-cell-sorting-10-k-1-standard-2-0-0. The human brain multiome sequencing data was downloaded from https://www.10xgenomics.com/cn/datasets/frozen-human-healthy-brain-tissue-3-k-1-standard-2-0-0. The human Colorectal scATAC-seq dataset was downloaded following the Signac Vignette at: https://github.com/stuart-lab/signac/blob/1.9.0/vignettes/mito.Rmd. The PBMC single-cell reference annotation was downloaded following the link in the Vignette at: https://github.com/stuart-lab/signac/blob/1.9.0/vignettes/pbmc_multiomic.Rmd.

## Acknowledgement

We thank Lionel Chiron and all the members of the Perié lab for helpful discussions. This work was funded by the European Research Council (ERC) under the European Union’s Horizon 2020 research and innovation programme ERC StG 758170-Microbar (to L.P.) and by the Institut Thématique Multi-Organismes (ITMO) Cancer of Aviesan within the framework of the 2021-2030 Cancer Control Strategy, administered by INSERM (to L.P.).

## Author contributions

Conceptualization was done by W.S., who also conducted formal analysis with the assistance of D.G. W.S. handled software development, data curation, and drafted the manuscript. L.P. secured funding, provided study supervision, and reviewed and edited the manuscript.

## Supplementary Figures

**Supplementary Figure 1.**
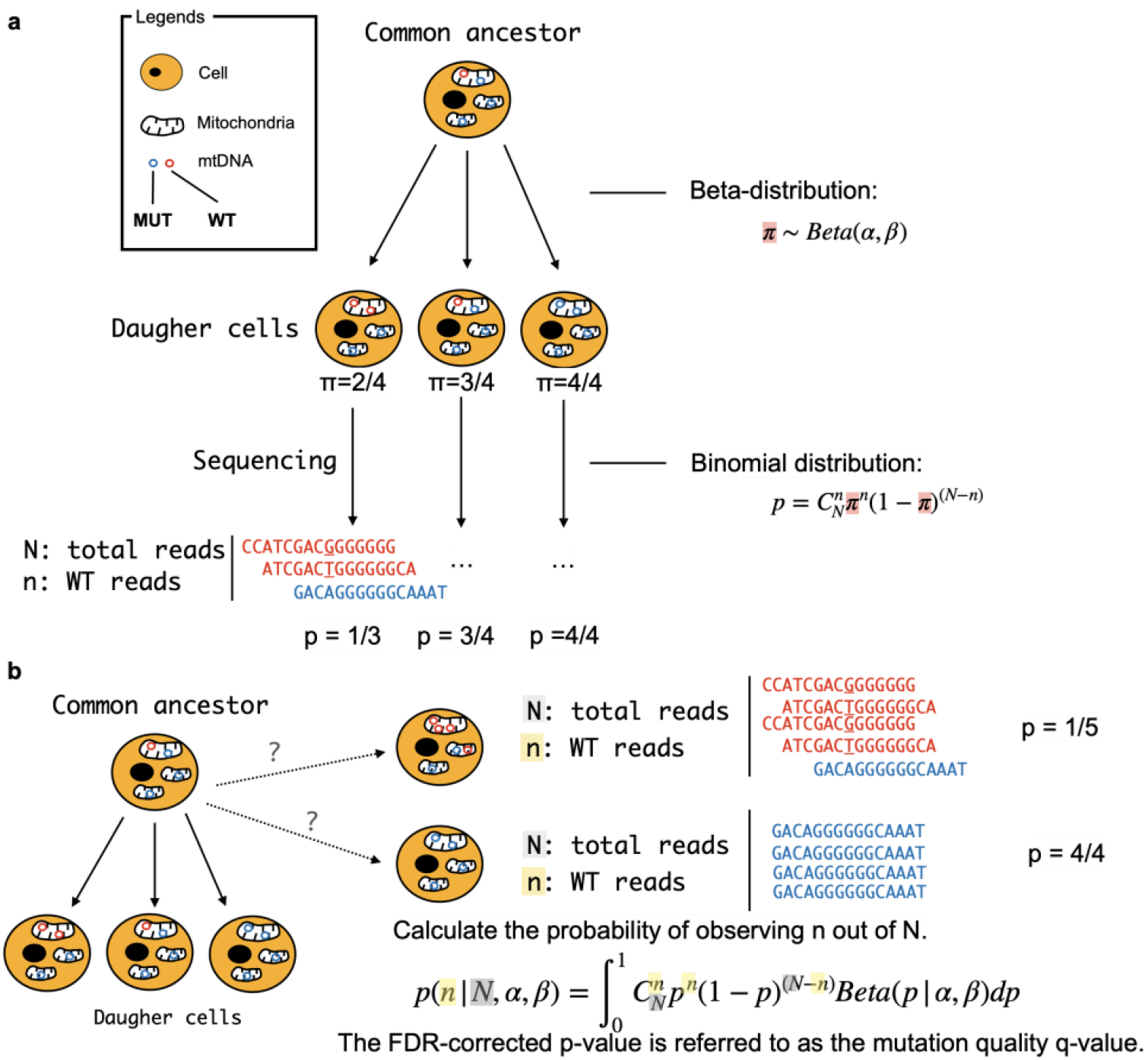
Identifying lineage informative mtDNA mutation using a beta-binomial model. (a) Modeling single-cell mtDNA allele frequency using a beta-binomial distribution as implemented into scMitoMut, it correspond to step 2 of Figure 1b. After sequencing, we obtained allele frequency by counting the reads. This frequency depends on the mtDNA heteroplasmy, mtDNA sampling, sequencing errors, and PCR errors and is modeled by a binomial process. The dynamic of mtDNA heteroplasmy ranges from 0 to 1 and depends on the dynamics of mtDNA as cells divide, this is modeled using a beta distribution. In the diagram, big yellow circles represent cells with black dot inside as nucleus. In each cell, there are multiple mitochondria and mtDNA. The small blue circles represent wild type mtDNA, while the red circles indicate mutant mtDNA. Both types are specific to a particular locus. (b) Assessing lineage-specific somatic mtDNA mutations per locus and per cell using the beta-binomial distribution, it corresponds to step 3 of figure 1b. If cells come from a shared ancestor and are wild-type, they should follow the beta-binomial distribution fitted on wild-type cells of step 2 of figure 1b, enabling us to calculate the probability of the cells being wild-type at a given locus, given its reads n at that locus and the total reads N. By adjusting the obtained probability with false discovery rate (FDR), the resulting adjusted probability acts as the mutation quality value, referred to as the q-value.

**Supplementary Figure 2.**
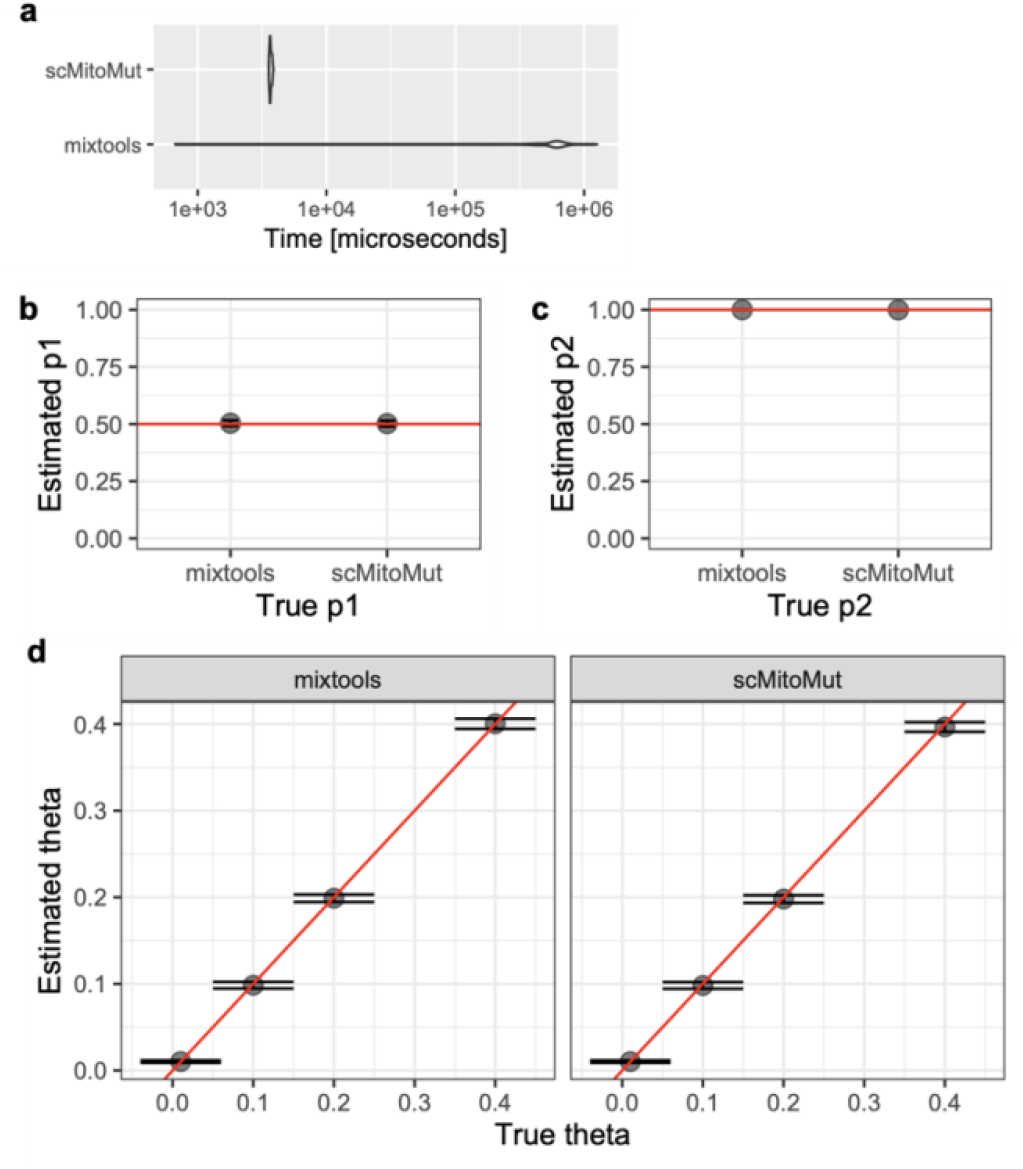
Performance of the binomial-mixture model in scMitoMut. **(a)** The time needed for model fitting of binomial-mixture model fitting for the R package mixtools and scMitoMut. The horizontal violin plot illustrates the distribution of time across 100 replications. The tested dataset is simulated binomial mixture distribution with two binomial distributions with *θ* = 0.*5, π*1 = 0.992 and *π*2 = 1 respectively. The n follows a log-normal distribution (log-mean = 2 and log-SD = 1) rounded up to the nearest non-negative integer. Each simulation includes 1000 observations. (b), (c) The fitting accuracy of parameters *π*_1_ (p1) and *π*_2_ (p2) obtained using the R package mixtools and scMitoMut. The red line indicates the true value. (c) The fitting accuracy for various values of parameter obtained using the R package mixtools and scMitoMut. The x and y axes represent the true and estimated values respectively, with the red line denoting the y=x diagonal. For fitting accuracy, the simulations follow this design: *π*_1_ *= 0*.*5, π*_2_ *=* 1, *θ =* 0.01, 0.1, 0.2, 0.4 and *n* following a log-normal distribution (log-mean=2, log-SD=1) rounded to the nearest non-negative integer. Each simulation contains 5000 observations. Error bars in (b)–(d) denote the mean ± SD.

**Supplementary Figure 3.**
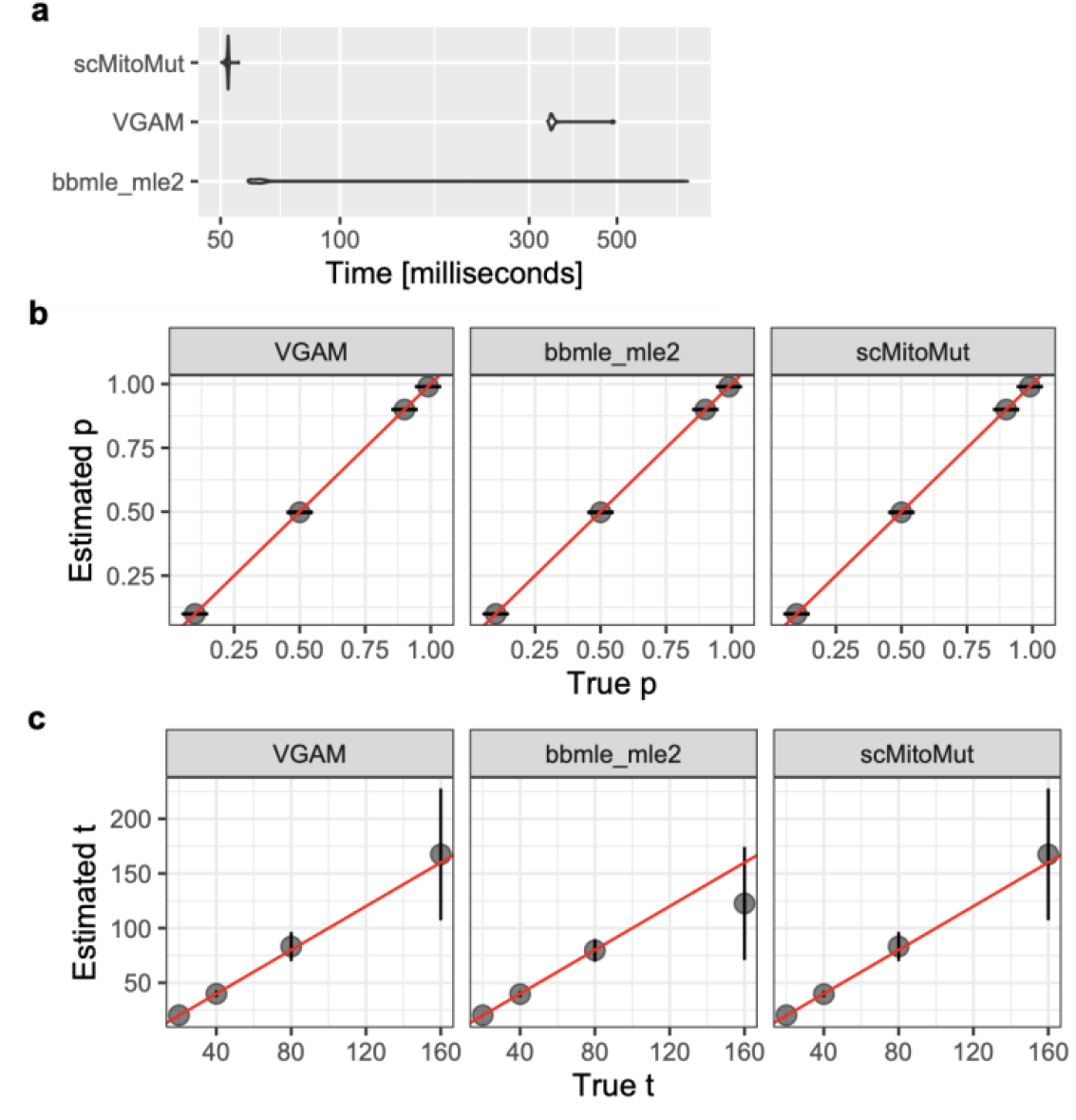
Performance of the beta-binomial model in scMitoMut. (a) The beta-binomial model fitting speed between the R package VGAM, bbmle and scMitoMut. The horizontal violin plot illustrates the distribution of time across 100 replications. The BBD dataset was simulated with a mean probability parameter of 0.5, and a super-dispersion parameter of 20. Each simulation includes 5000 observations. (b), (c) The fitting accuracy for parameter *θ* and *ϕ* using the R packages VGAM, bbmle, and scMitoMut. The red line denotes y = x diagonal. The parameter values are: mean probability parameter *θ* of 0.1, 0.5, 0.9, and 0.99; dispersion parameter *ϕ* of 20, 40, 80, 160. In the simulation, *n* adheres to a log-normal distribution with a log-mean of 2 and log-SD of 1, subsequently rounded up to the nearest non-negative integer. Each simulation encompasses 5000 observations. The error bars in panels B and C represent the mean ± SD.

**Supplementary Figure 4.**
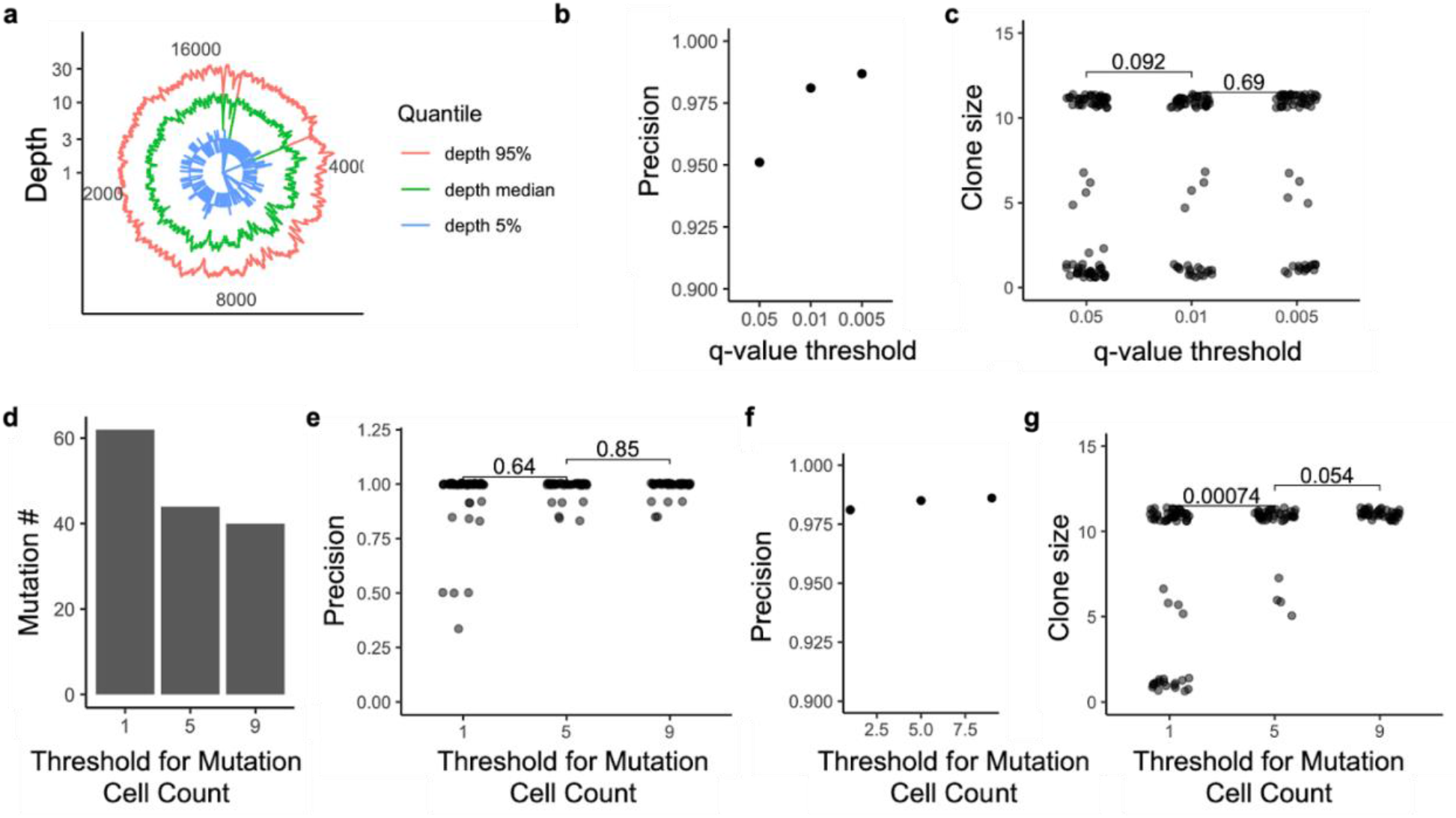
Effect of q-value and number of mutant cells on the mutation calling using scMitoMut on single cell genome sequencing of cell line mixture dataset. (a) Polar plot displaying the quantile of mtDNA sequencing depth in the 10X Genomics single-cell genome sequencing dataset. Mitochondrial locus is shown by angle and sequence depth by radius. Blue, green, and pink indicate the 5th, 50th, and 95th percentiles, respectively, of depth across the population. All cells were analysed with no filtering. (b) Precision of mutation calling with different q-value thresholds. Precision is calculated by dividing the number of MKN45 mutant cells by the total number of mutant cells, corresponding to a clone size weighted average precision per mutation. (c)The number of cells per mutation (clone size) for different q-value thresholds. Each dot represents a mutation, Only mutations with at least one MKN45 cell passing the threshold were included in the analysis. The two-sided Wilcoxon test was used to compare mutant cell numbers between groups, with p-values displayed on top. The n is 79, 62 and 59 for q-value threshold of 0.05, 0.01 and 0.005 respectively. (d) Number of mutations detected with different thresholds of mutant cell numbers. Mutant cells were first filtered using a q-value <0.01. (e) Precision of mutations with different thresholds of mutant cell numbers. Each dot represents a mutation; mutant cells were first filtered using a q-value < 0.01. Precision was defined as in (b). The two-sided Wilcoxon Test was used to compare mutation precision, with the *p*-value displayed at the top. The n is 64, 44 and 40 for mutation cell count threshold of 1, 5 and 9 respectively. (f) Precision of mutation calling with different thresholds of mutant cell numbers. Precision is calculated by dividing the number of MKN45 mutant cells by the total number of mutant cells, corresponding to a clone size weighted average precision per mutation. (g) Number of cells per mutation for different mutant cell number thresholds. Similar to (c), each dot represents a mutation. Statistical analysis was performed using the two-sided Wilcoxon test, with *p*-values indicated in the figure. The n is 64, 44 and 40 for mutation cell count threshold of 1, 5 and 9 respectively.

**Supplementary Figure 5.**
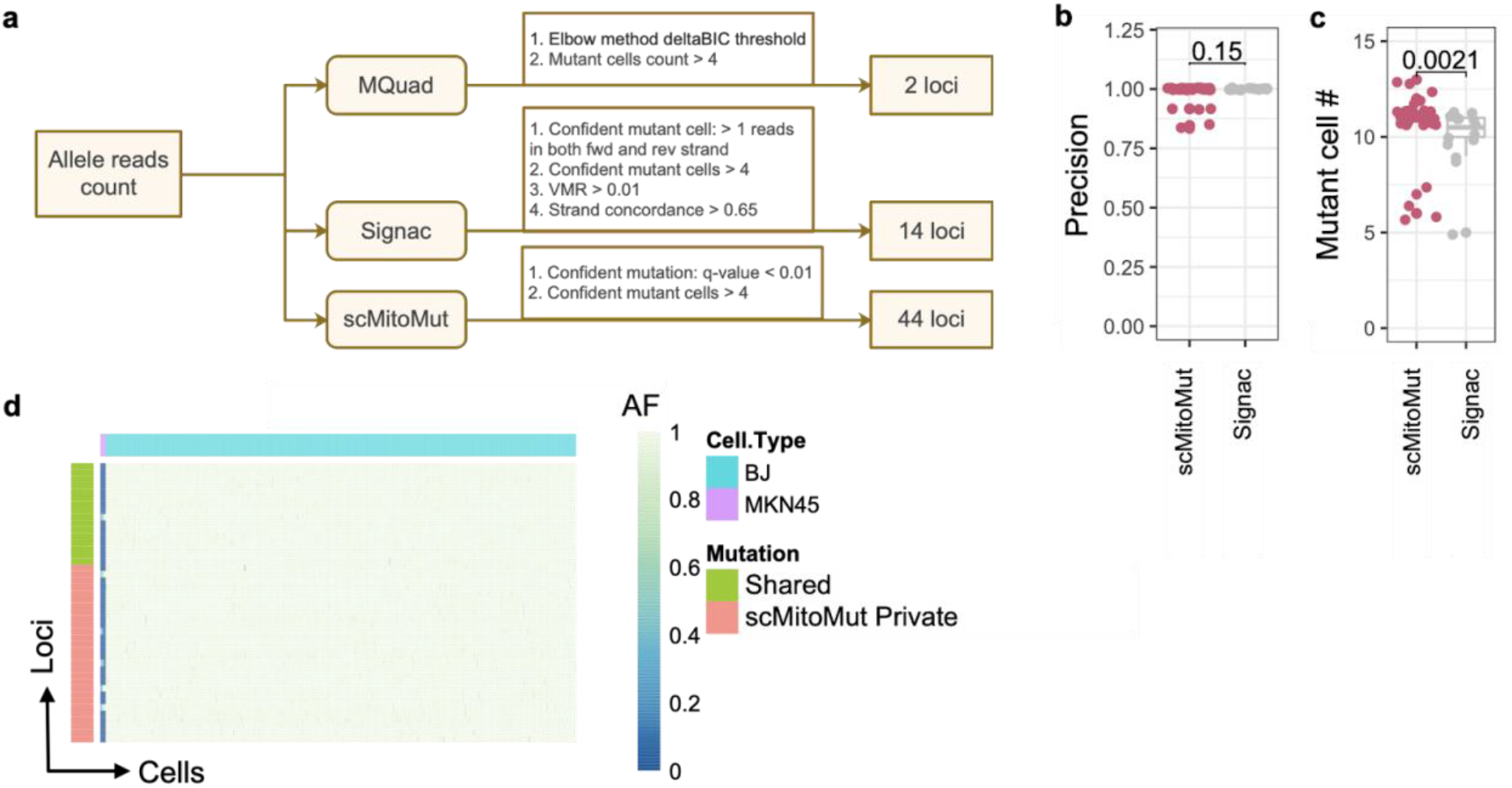
Benchmarking scMitoMut over SOTA with single cell genome sequencing of cell line mixture dataset. (a) Parameters used for scMitoMut, Signac and MQuad. For scMitoMut, mutant cells are identified when the q-value < 0.01 and mutant cells >= 5. In the Signac pipeline, we followed default settings in its vignette: the informative locus should have a variant mean ratio (VMR) > 0.1; strand correlation > 0.65; confident mutant cells >= 5 (confident mutant cell is defined by having more than two mutant reads in forward and reverse strand reads respectively). For MQuad, we used the auto-determined delta Bayesian Information Criterion (deltaBIC) threshold to filter mutations. (b) The scatter plot shows the lineage precision of scMitoMut and Signac results. The mutations with at least one MKN45 cell were analysed. Each dot in the scatter plot represents a mutation; Precision was calculated as the ratio of mutant MKN45 to total mutant cells for each mutation. Statistical analysis was conducted using the two-sided Wilcoxon Test, with p-values indicated in the plot. n = 14 mutations in scMitoMut and n = 9 mutations in Signac. (c) The scatter plot showing the mutant cell number for each mutation with scMitoMut and Signac. The mutations with at least one MKN45 cell are analysed. Statistical analysis was conducted using the two-sided Wilcoxon test, with *p*-values indicated in the plot. n = 14 mutations in scMitoMut and n = 9 mutations in Signac. (d) The mutation heatmap shows allele frequency of mutations per cell (columns) and per locus (rows). Blue shading represents wild type allele frequency. An annotation bar above the heatmap indicates the cell type (purple for MKN45 cells, and blue for BJ cells). Another annotation bar on the right shows mutations exclusively identified by scMitoMut, in red, and shared mutations between scMitoMut with MQuad and Signac, in green.

**Supplementary Figure 6.**
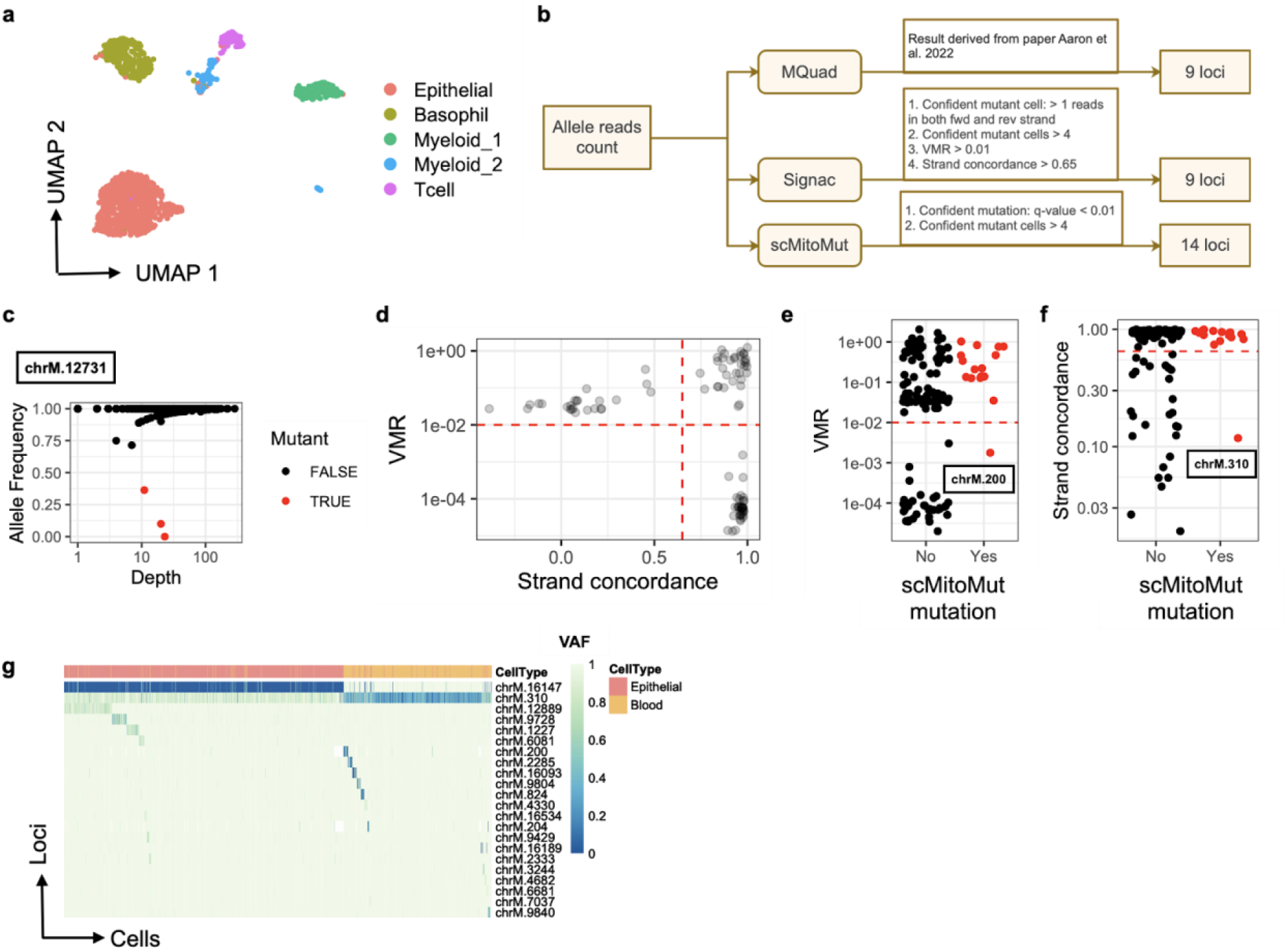
scMitoMut separates epithelial and blood lineage in a CRC dataset. (a) UMAP representation of immune and cancer epithelial cells from the CRC dataset, colored by cell type using Seurat with default parameters. (b) Parameters used for scMitoMut, Signac and MQuad. For scMitoMut, mutant cells are called if the q-value < 0.01 and mutant cells >= 5. In the Signac pipeline, we followed default settings in its vignette: the informative locus should have a Variant Mean Ratio (VMR) > 0.1; strand correlation > 0.65; confident mutant cells >= 5 (confident mutant cell is defined by having more than two mutant reads in forward and reverse strand reads respectively). For MQuad, we used the auto-determined delta Bayesian Information Criterion (deltaBIC) threshold to filter mutations. (c) Sequencing depth and allele frequency of locus chrM.12731. Each dot represents a cell, with cells having scMitoMut mutation q-value < 0.01 highlighted in red. (d) Variant mean ratio (VMR) and strand correlation distribution from the Signac result. Each dot is a mutation. Only mutations with at least 5 confident mutant cells are included in the plot. Each dot represents a mutation locus, and red lines indicate the default thresholds in Signac for VMR (0.01) and strand correlation (0.65). (e) and (f) Signac VMR or strand concordance profiling with scMitoMut mutations called by: q-value < 0.05 and at least 5 mutant cells. Each dot is a locus. The x-axis shows the mutation status identified by scMitoMut, and the y-axis is the VMR value of the mutation (e) or strand concordance of mutation reads (f). The red line represents the threshold used by Signac which is 0.01 for VMR 0.01 and 0.65 for strand concordance 0.65. (g) The mutation heatmap showing the mutation allele frequency per cell (columns) and per mutation (rows). The shading of blue within the heatmap represents the wild type allele frequency. An annotation bar above the heatmap indicates the cell type with yellow of blood and red of cancer epithelial cells.

**Supplementary Figure 7.**
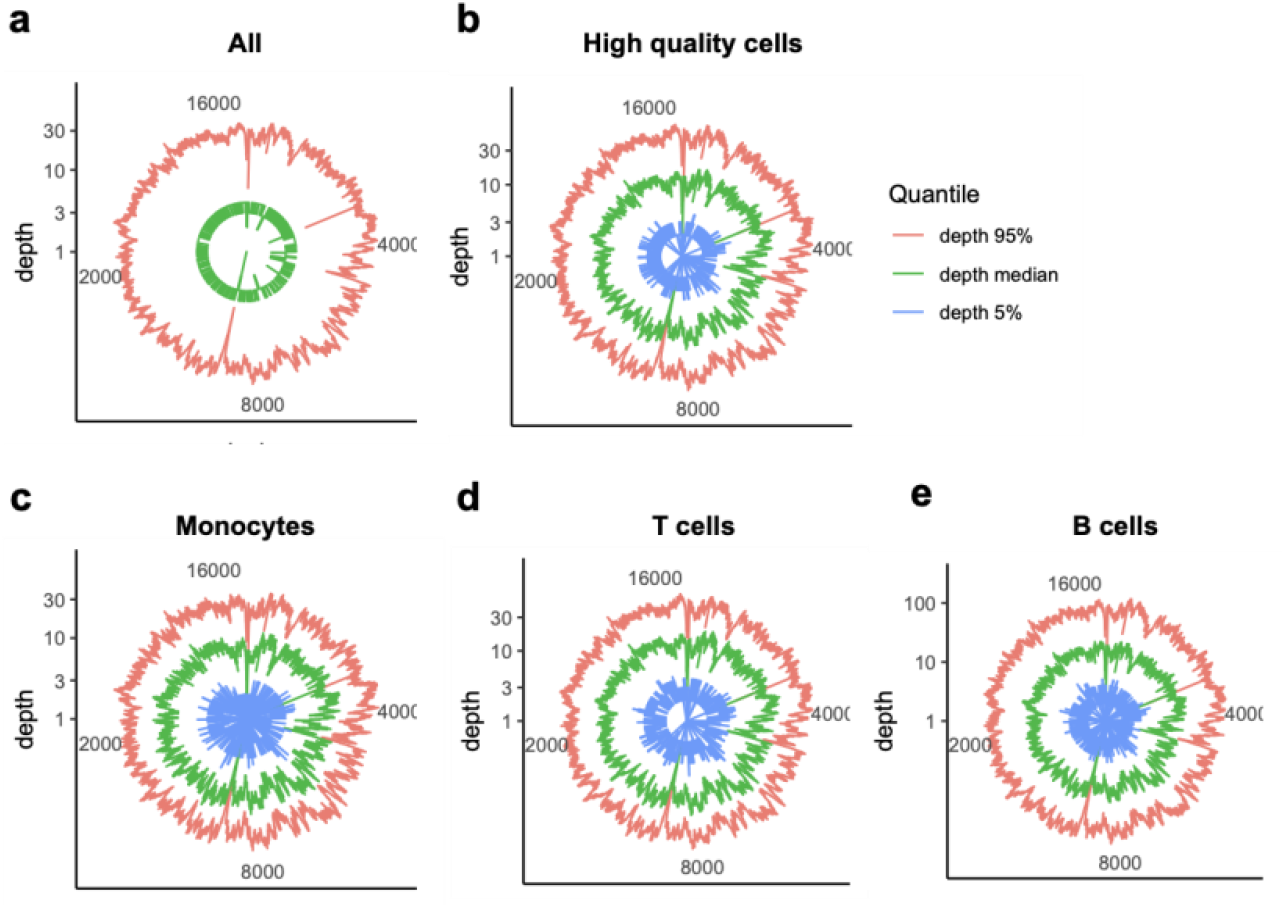
scMitoMut separates epithelial and blood lineage in 10X PBMC dataset. (a) Polar plot displaying the quantile of mtDNA sequencing in the 10X PBMC multiome dataset scATACSeq unfiltered cells. Mitochondrial locus is shown by angle and sequence depth by radius. Blue, green, and pink indicate the 5th, 50th, and 95th percentiles, respectively, of depth across the population. (b) Same as (a) but for high quality cells, which have a median mtDNA sequencing depth greater than 5. (c-e) Same as (a) but for monocytes (c), T cells (d) and B cells (e) mtDNA sequencing depths among the high-quality cells of (b).

**Supplementary Figure 8.**
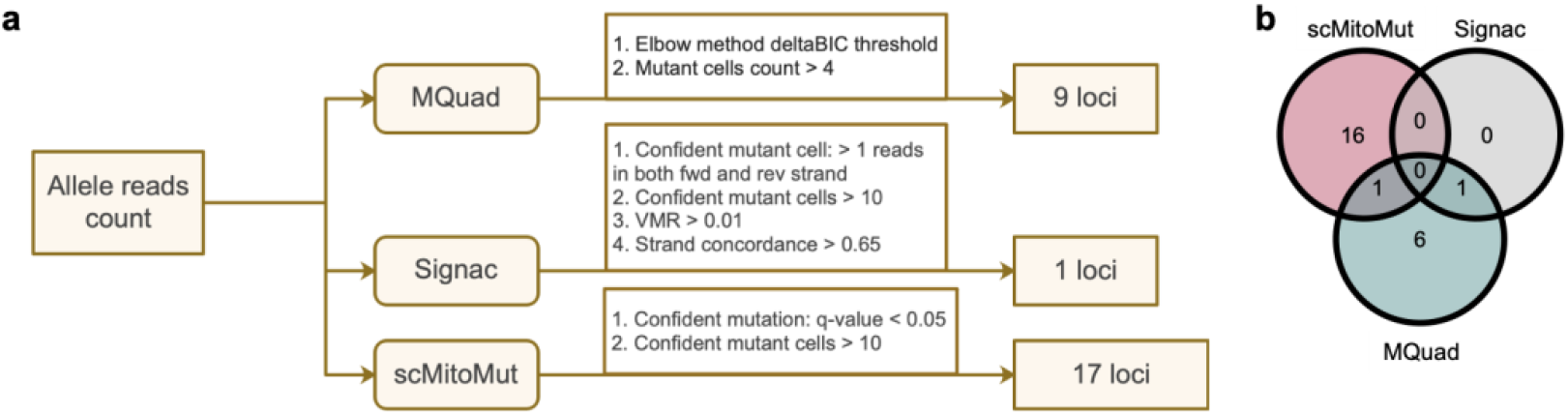
Benchmarking scMitoMut over SOTA methods using the PBMC 10X multiome dataset. (a) Parameters used for scMitoMut, Signac and MQuad. For scMitoMut, mutant cells are identified when the q-value < 0.01 and mutant cells > 10. In the Signac pipeline, we followed default settings in its vignette: the informative locus should have a Variant Mean Ratio (VMR) > 0.1; strand correlation > 0.65; confident mutant cells >= 5 (confident mutant cell is defined by having more than two mutant reads in forward and reverse strand reads respectively). For MQuad, we used the auto-determined delta Bayesian Information Criterion (deltaBIC) threshold to filter mutations. (b) Number of mutations identified by the three tools: scMitoMut in red, Signac in gray, and MQuad in green using the parameters in (a).

**Supplementary Figure 9.**
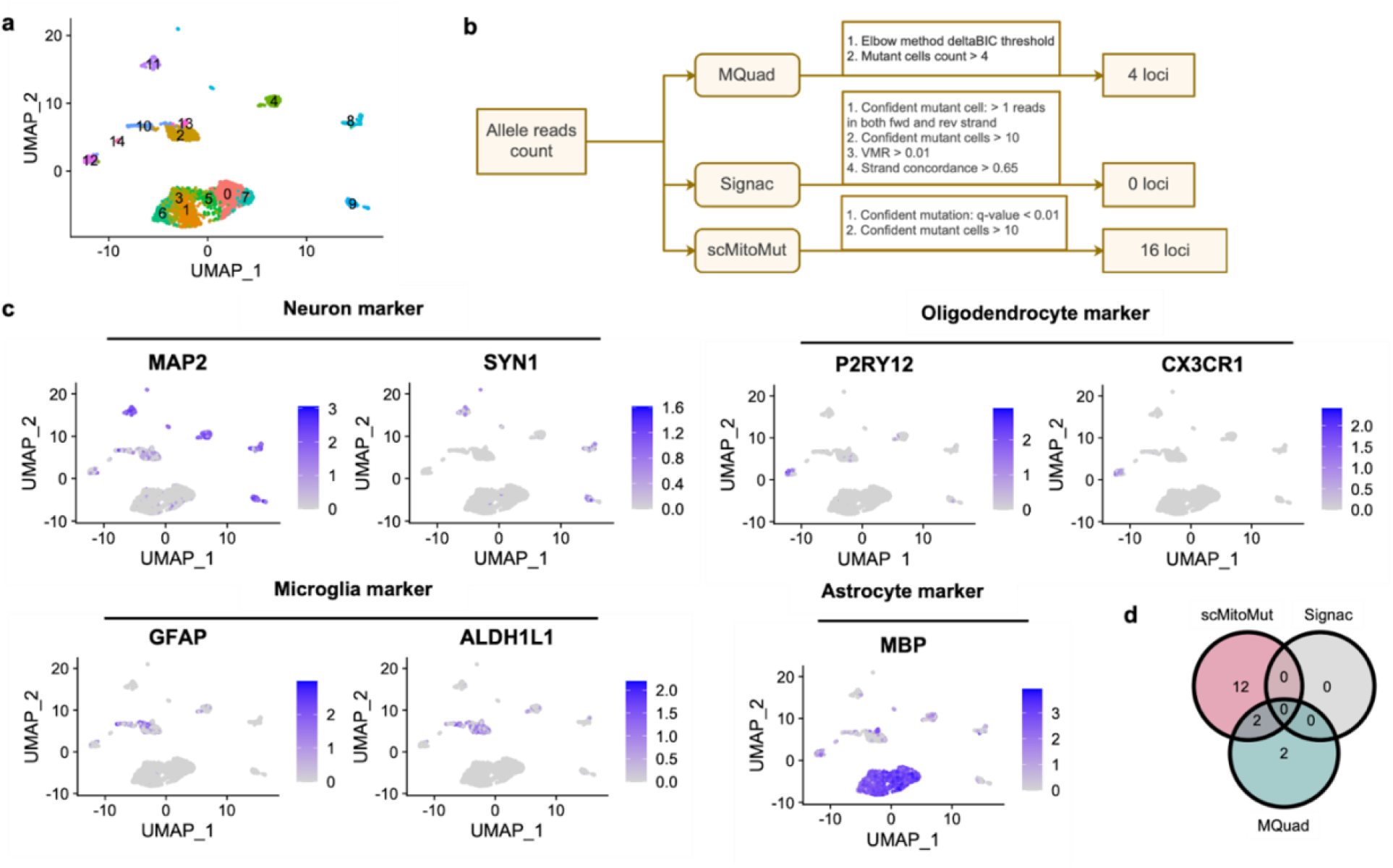
scMitoMut reveals lineage informative mitochondrial mutations in 10x Brain multiome dataset. (a) UMAP of the brain 10X multiome dataset. The clusters identified by RNA are labeled by cluster id and annotated by colors using Seurat with default parameters. (b) Parameters used for scMitoMut, Signac and MQuad. For scMitoMut, mutant cells are identified with the q-value < 0.01 and mutant cells > 10. In the Signac pipeline, we followed default settings in its vignette: the informative locus should have a Variant Mean Ratio (VMR) > 0.1; strand correlation > 0.65; confident mutant cells >= 5 (confident mutant cell is defined by having more than two mutant reads in forward and reverse strand reads respectively). For MQuad, we used the auto-determined delta Bayesian Information Criterion (deltaBIC) threshold to filter mutations. (c) same UMAP as in (a) but color-coded for the expression of gene markers of the neuron, astrocyte, microglia and oligodendrocyte. (d) Number of mutations identified by the three tools: scMitoMut in red, Signac in gray, and MQuad in green using the parameter in (b).

