## supplementary methods for "scMitoMut for calling mitochondrial lineage–related mutations in single cells"

### Fit binomial mixture model with Expectation maximization (EM) algorithm

For a binomial mixture model (1) with 2 component $k = [1,2]$ which represent two status: wild type and mutant. The binomial mixture model has parameters $\Theta = [\pi_1, \theta_1, \pi_2, \theta_2]$.

$$P(n_i|N_i, \Theta) = \pi_1 Binom(n_i|N_i, \theta_1) + \pi_2 Binom(n_i|N_i, \theta_2) \qquad (1)$$

We fit the model by expectation maximization algorithm.

**Initialization**: Start by randomly initializing the parameters of the two binomial distributions $\Theta$, and assign it to $\Theta_{old}$.

**Expectation step (E-step)**: Given the current parameters $\Theta_{old}$, calculate the probability (2) that each cell come from distribution $k$.

$$P(z_i = k|n_i, \Theta_{old}) = [\Sigma_{l=1}^{2} \frac{\pi_{1,old}}{\pi_{k,old}} (\frac{\theta_{1,old}}{\theta_{k,old}})^{n_i} (\frac{1-\theta_{1,old}}{1-\theta_{k,old}})^{N_i - n_i}]^{-1} \qquad (2)$$

**Maximization step (M-step)**: Update the parameters to have new $\Theta$.

$$\pi_k = \frac{1}{S} \Sigma_{i=1}^{S} P(z_i = k|n_i, \Theta_{old}) \qquad (3)$$

$$\theta_k = \frac{\Sigma_{i=1}^{S} n_i P(z_i = k|n_i, \Theta_{old})}{\Sigma_{j=1}^{S} N_j P(z_j = k|n_j, \Theta_{old})} \qquad (4)$$

In (3) and (4), $S$ is the cell number.

**Iteration**: Assign the updated $\Theta$ to $\Theta_{old}$, and repeat E-step and M-step until the log likelihood function (5) converges.

$$\begin{aligned} lnf(\vec{n}, \vec{N}) &= \Sigma_{i=1}^{S} lnP(n_i|N_i, \Theta) \\ &= \Sigma_{i=1}^{S} ln(\pi_1 Binom(n_i|N_i, \theta_1) + \pi_2 Binom(n_i|N_i, \theta_2)) \qquad (5) \end{aligned}$$

### Fit beta binomial distribution with Maximum Likelihood Estimation (MLE) algorithm

For the beta-binomial distribution (6) with parameter $\alpha$ and $\beta$.

$$P(n_i|N_i,\alpha,\beta) = \binom{N_i}{n_i}\frac{\Gamma(\alpha+\beta)}{\Gamma(\alpha)\Gamma\beta}\Gamma(\alpha+n_i)\frac{\Gamma(\beta+N_i-n_i)}{\Gamma(\alpha+\beta+N_i)} \qquad (6)$$

We fit it by MLE algorithm using the allele count data from $S$ cells.

Firstly, we define the likelihood (7) of the parameters for observed allele counts.

$$f(\vec{n},\vec{N}) = \Pi_{i=1}^{S}(\binom{N_i}{n_i}\frac{\Gamma(\alpha+\beta)}{\Gamma\alpha\Gamma\beta}\Gamma(\alpha+x_i)\frac{\Gamma(\beta+n_i-x_i))}{\Gamma(\alpha+\beta+n_i)} \qquad (7)$$

From (7), derived the log likelihood function (8).

$$\begin{aligned} lnf(\vec{n},\vec{N}) &= \Sigma_{i=1}^{S} ln(\binom{N_i}{n_i}\frac{\Gamma(\alpha+\beta)}{\Gamma\alpha\Gamma\beta\Gamma(\alpha+\beta+N_i)}\Gamma(\alpha+n_i)\Gamma(\beta+N_i-n_i)) \\ &= S[ln\Gamma(\alpha+\beta) - ln\Gamma(\alpha) - ln\Gamma(\beta)] - \Sigma_{i=1}^{S} ln\Gamma(\alpha+\beta+N_i) + \Sigma_{i=1}^{S} ln\binom{N_i}{n_i} \\ &+ \Sigma_{i=1}^{S} ln\Gamma(\alpha+n_i) + \Sigma_{i=1}^{S} ln\Gamma(\beta+N_i-n_i) \qquad (8) \end{aligned}$$

After beeing initialized, the parameter $\alpha$ and $\beta$ are fitted using Newton-Raphson method by maximizing the likelihood.

$$[\alpha\ \beta\ ]_{New} = [\alpha\ \beta\ ]_{old} - J^{-1}([\alpha\ \beta\ ]_{old})\nabla f([\alpha\ \beta\ ]_{old}) \qquad (9)$$

In equation (9), the Jacobian matrix is defined as (10).

$$J = \left[\frac{\partial}{\partial\alpha^2}lnf(\vec{n},\vec{N})\ \ \frac{\partial}{\partial\alpha\partial\beta}lnf(\vec{n},\vec{N})\ \ \frac{\partial}{\partial\alpha\partial\beta}lnf(\vec{n},\vec{N})\ \ \frac{\partial}{\partial\beta^2}lnf(\vec{n},\vec{N})\right] \qquad (10)$$

The equations (11), (12), (13), (14), and (15) define the partial derivatives with respect to $\alpha$ and/or $\beta$ in the log-likelihood function, that are used in equation (9) and (10).

$$\frac{\partial}{\partial\alpha}lnf(\vec{n},\vec{N}) = S[\gamma_0(\alpha+\beta) - \gamma_0(\alpha)] - \Sigma_{i=1}^{S}\gamma_0(\alpha+\beta+N_i) + \Sigma_{i=1}^{S}\gamma_0(\alpha+n_i) \qquad (11)$$

$$\frac{\partial}{\partial\beta}lnf(\vec{n},\vec{N}) = S[\gamma_0(\alpha+\beta) - \gamma_0(\beta)] - \Sigma_{i=1}^{S}\gamma_0(\alpha+\beta+N_i) + \Sigma_{i=1}^{S}\gamma_0(\beta+N_i-n_i) \qquad (12)$$

$$\frac{\partial}{\partial \alpha^2} lnf(\vec{n}, \vec{N}) = S[\gamma_1(\alpha + \beta) - \gamma_1(\alpha)] - \Sigma_{i=1}^{S} \gamma_1(\alpha + \beta + N_i) + \Sigma_{i=1}^{S} \gamma_1(\alpha + n_1) \qquad (13)$$

$$\frac{\partial}{\partial \beta^2} lnf(\vec{n}, \vec{N}) = S[\gamma_1(\alpha + \beta) - \gamma_1(\beta)] - \Sigma_{i=1}^{S} \gamma_1(\alpha + \beta + N_i) + \Sigma_{i=1}^{S} \gamma_1(\beta + N_i - n_i) \qquad (14)$$

$$\frac{\partial}{\partial \alpha \partial \beta} lnf(\vec{n}, \vec{N}) = N\gamma_1(\alpha + \beta) - \Sigma_{i=1}^{N} \gamma_1(\alpha + \beta + n_i) \qquad (15)$$
